# Learning reweights the decision dynamics of cortico-basal ganglia-thalamic pathways from deliberation to commitment

**DOI:** 10.64898/2026.02.17.706272

**Authors:** Zhuojun Yu, Jonathan E. Rubin, Timothy Verstynen

**Affiliations:** Department of Psychology & Neuroscience Institute, Carnegie Mellon University, Pittsburgh, Pennsylvania, United States of America; Department of Mathematics, University of Pittsburgh, Pittsburgh, Pennsylvania, United States of America; Center for the Neural Basis of Cognition, Pittsburgh, Pennsylvania, United States of America

## Abstract

Mammals flexibly adjust their decision strategies in dynamic environments based on prior experience. The cortico-basal ganglia-thalamic (CBGT) circuit is recognized as a critical driver of this adaptability, yet how plasticity-induced modifications to CBGT dynamics translate into modifications of decision policies remains poorly understood. Here we simulate learning in a biologically-grounded spiking CBGT model that learns to select a rewarded action via dopamine-dependent plasticity at corticostriatal synapses. Relying on three previously identified control ensembles (responsiveness, pliancy, and choice) within CBGT circuits, we disentangle the distinct roles these subnetworks have in reshaping decision trajectories across learning. Control ensemble dynamics were mapped onto the evolution of evidence accumulation, revealing within-trial parameter adjustments that shape decision dynamics and outcomes. Our results emphasize that learning optimizes not only what choice is favored, but also how the phases within the decision unfold. Early in the decision, learning accelerates evidence accumulation by driving the activity of corticothalamic and direct pathways. During later deliberation, this drive is temporarily restrained, and the indirect and pallidostriatal pathways become more critical for maintaining decision thresholds, preventing premature commitment despite the increased choice bias. As the system approaches the decision point, the direct pathway regains dominance, triggering boundary collapse to facilitate action selection. This mechanism effectively shifts decisions from deliberative to committed reward-directed choices, improving both speed and accuracy while preserving system stability and control throughout the process.

**Author summary:** We investigate how specific subnetworks within cortico-basal ganglia-thalamic (CBGT) circuits reshape decision-making through dopamine-dependent plasticity at corticostriatal synapses. Using simulations of a simple two-choice task with a single rewarded target, we identify phase-specific reconfigurations of subnetwork activity that emerge over the course of learning. We then link these circuit-level changes to an algorithmic description of behavior by mapping them onto decision policies within a dynamic evidence accumulation framework. This analysis reveals a learning-driven shift from deliberative to more committed decision strategies, achieved through flexible coordination of decision boundaries and evidence accumulation rates. Together, these adjustments optimize decision speed and accuracy while preserving caution during ongoing choices. Our findings provide mechanistic insight into how plasticity within CBGT circuits supports adaptive decision-making across different levels of uncertainty and task demands.

## Introduction

Behavioral adaptability is critical for survival in dynamic or uncertain environments. To meet changing conditions and task demands, animals must continually integrate both internal and external cues to select appropriate actions. This adaptive action selection relies heavily on plasticity within the cortico-basal ganglia-thalamic (CBGT) circuit, which can shape the evidence accumulation process during decision-making [1–5]. A critical target for plasticity within the CBGT circuit is the corticostriatal synapses, who receive dopaminergic signals that modify how cortical input drives CBGT dynamics and subsequent behavior [6–10].

Dopaminergic modulation of the corticostriatal synapses can alter the overall CBGT dynamics via at least two structurally and functionally dissociable routes: the *direct* and *indirect* pathways [11, 12]. Cortical inputs to the direct pathway result in inhibition of basal ganglia outputs and disinhibition of downstream thalamic sites, whereas activation of the indirect pathway induces opposite effects. These pathways originate in the striatum with two classes of spiny projection neurons (SPNs) classified according to their predominant dopamine receptor type. D1-expressing SPNs (dSPNs) start the direct pathway (Fig 1A, green) and exhibit synaptic potentiation when driven by cortical inputs in the presence of positive phasic dopamine signals. D2-expressing SPNs (iSPNs), by contrast, start the indirect pathway (blue), and their synapses are depressed by cortical inputs along with positive phasic dopamine signals. These opposing plasticity effects on the competing direct and indirect pathways are believed to regulate dynamic processing of cortical activity by the basal ganglia [5, 13].

**Fig 1.**
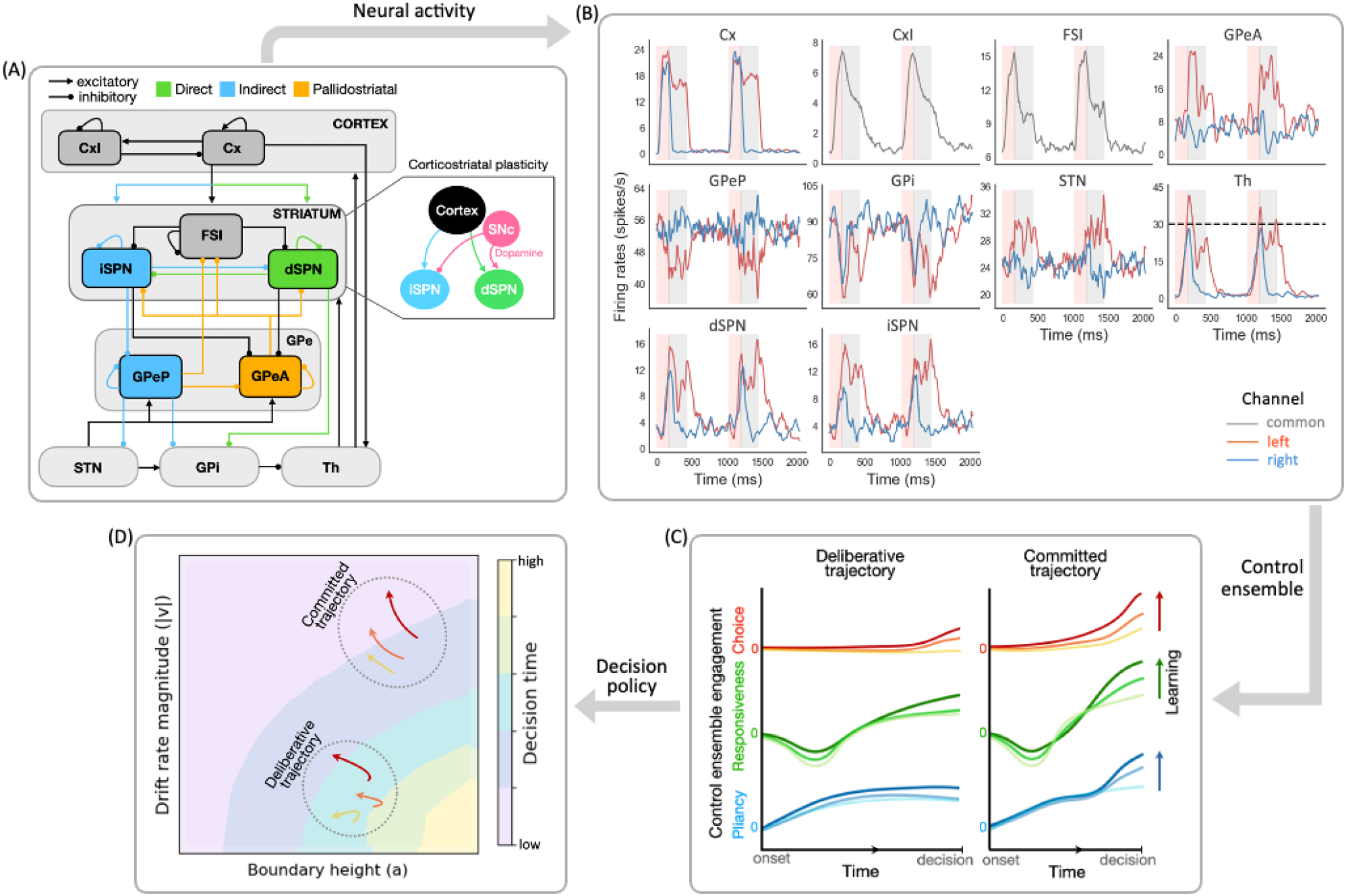
Steps in analyzing learning effects on decision-making within CBGT circuits. (A) The CBGT circuit, with connections color-coded as follows: classical direct pathway in green, classical indirect pathway in blue, and pallidostriatal pathway in gold. Note that the colors highlight the key nuclei shaping each pathway’s influence, rather than every anatomical component. For example, internal globus pallidus (GPi) participates in both direct and indirect pathways, and subthalamic nucleus (STN) is part of the indirect pathway, but we color only the nuclei whose activity most clearly reflects each pathway’s role during decisions. Arrows ending in dots indicate postsynaptic sites of inhibitory connections, while arrows ending in triangles indicate excitatory connections. Plasticity changes at corticostriatal synapses are induced by dopaminergic signals associated with rewards following decisions (pink arrows). Cx, cortical neurons; CxI, inhibitory cortical interneurons; FSI, fast spiking interneurons; dSPN, direct spiny projection neurons; iSPN, indirect spiny projection neurons; GPe, external globus pallidus; GPeP, prototypical neurons; GPeA, arkypallidal neurons; GPi, internal globus pallidus; STN, subthalamic nucleus; Th, thalamus; SNc, substantia nigra pars compacta. (B) Example of firing rate time courses for all CBGT nuclei in a two-choice task. Each red (blue) trace corresponds to activity in the left (right) action channel in a CBGT region, and the grey traces correspond to the populations (i.e., CxI and FSI) common to both action channels. Pink regions represent the decision phase, occurring before the thalamus of one of the action channels reaches the decision threshold of 30 Hz (dashed black line in Th panel). Grey regions represent the consolidation phase, where partial cortical input to the selected channel is sustained. The unshaded regions represent the inter-trial interval. (C) Time courses of control ensemble engagement, including choice (red), responsiveness (green), and pliancy (blue), over the decision phase for two decision types, deliberative and committed, mapped from the firing rate time courses. Color intensity increases with learning progression. (D) Trajectories of DDM parameters (drift rate and boundary height), mapped from the time courses of control ensemble engagement, showing how the decision policy shifts with learning. Background manifolds indicate decision times derived from DDM simulations, with yellow (purple) colors representing long (short) decision times.

There are several other characteristics of the canonical CBGT model that are important for understanding how these circuits regulate behavioral adaptability. First, CBGT circuits are thought to be spatiotemporally organized into discrete action representations, termed *action channels* [14–17], each comprising both direct and indirect pathway components, with the former facilitating and the latter suppressing selection of the represented action. In addition, the dynamics of direct and indirect pathway activity are influenced by bidirectional control signals emanating from the external segment of the globus pallidus (GPe; see [18, 19]). The GPe is now seen as being composed of two major classes of neurons, namely prototypic (GPeP), projecting towards basal ganglia output nuclei, and arkypallidal (GPeA), which project back to the striatum and influence both iSPNs and dSPNs [18, 20–22]. Through these pallidostriatal feedback projections, GPeA cells regulate striatal activity in contexts requiring reactive inhibitory control [23–25]. Thus, GPeA neurons and their projections comprise a third major pathway in recent updates of the canonical CBGT circuit—the *pallidostriatal* pathway (Fig 1A, gold). By including these three essential CBGT pathways in a computational model, our work aims to explain how dopaminergic plasticity results in changes in the CBGT circuit dynamics that unfold during the course of individual decisions, revealing how feedback learning promotes adaptive decision-making.

Prior computational work [26–29] has demonstrated mappings between the circuit-level dynamics of CBGT pathways and algorithmic representations of the evidence accumulation process, often captured by frameworks like the drift-diffusion model (DDM; see [30]). Our contribution in this space has identified three distinct CBGT subnetworks, or *control ensembles*, whose engagement translates into a specific influence on DDM parameters [29, 31, 32]. In this way the control ensembles can be thought of as dynamic subnetworks that have distinct impacts on the selection of actions. Corticostriatal plasticity can shift the overall engagement of control ensembles, leading to changes in the emergent decision policies in response to learned contingencies [31]. However, this work assumed time-invariant decision policy parameters, at least within the time course of a single decision. Thus it did not account for the fact that decision-makers can adjust their instantaneous strategies as a decision unfolds in response to real-time sensory feedback changes. Consistent with this idea, there is a growing body of work [33–35] exploring models of dynamic evidence accumulation, in which certain parameters, particularly the rate of evidence accumulation (drift rate) and level of evidence needed to make a decision (boundary height), are allowed to vary within a trial, to more realistically represent evolving decision policies [32]. Yet, at this point, it remains unclear how plasticity-driven changes in the temporal dynamics of CBGT control ensembles give rise to rapid, *within-trial* variations in decision policies.

We recently proposed a novel computational framework [32], called CLAW (Circuit Logic Assessed via Walks), for tracking the moment-to-moment propagation of neural activity as it progresses through CBGT circuits in the context of a simple two-choice task. The CLAW analysis revealed that, in the absence of learning, the complex interactions of CBGT pathways contribute to three functional phases of a decision: *launching, deliberation*, and *commitment*. The way in which trial trajectories progress through these phases leads to distinct decision types (scenarios). The CLAW analysis gives us a tool by which to understand the dynamic reconfiguration of decision policies as a decision unfolds.

Here we use the CLAW analysis to understand how dopamine-dependent plasticity at corticostriatal synapses shifts the instantaneous dynamics of the subsequent process of evidence accumulation up to a decision threshold. We first set out to replicate our previous findings of the three functional phases of a decision. Next, we proceed to convert the firing rate activity of CBGT nuclei into the temporal evolution of control ensemble engagement, as sketched in Fig 1B-C. This translation enables us to uncover the underlying CBGT substrates responsible for driving shifts in the distribution of decision types and their associated outcomes as learning advances. From there we establish a mapping from the temporal evolution of control ensembles to the DDM space (Fig 1C-D), revealing how boundary height and drift rate change as trajectories proceed through different decision phases. This mapping helps us to characterize how the resulting decision strategies are dynamically tuned through trial-by-trial learning. Progressing from biologically detailed mechanisms to algorithmically abstract representations, our results show that by reshaping the engagement of specific CBGT subnetworks, plasticity drives targeted adjustments in decision-related policies, steering decisions from deliberation to reward-aligned commitment while coordinating speed, control, and choice accuracy.

## Results

### Distinct decision types arise from CBGT dynamics

Our primary goal was to capture how the dynamics of CBGT circuits, during the evidence accumulation process, can evolve with learning over time. To this end we simulated 100 instances of a spiking computational model of CBGT networks [36] performing a simple two-choice task. Each network (reproduced from the “slow” networks in our past work [32]) featured a distinct configuration of synaptic weights that produced firing rates of all CBGT populations within experimentally observed ranges (see also [29, 31]). In the absence of plasticity these networks were unbiased with respect to left and right choices, as both action channels shared identical connectivity parameters and the evidence for the two options, represented by the input current to the two cortical populations, was always equivalent. This design ensured that changes in decision outcomes across different conditions resulted solely from learning-induced plasticity in the network.

To investigate how learning progressively tunes network dynamics, we began by simulating 50 trials per network with plasticity off, serving as a baseline prior to learning. Learning was then introduced in the form of a deterministic reward that was consistently assigned to the left choice. The magnitude of the phasic dopamine response following each trial was based on the reward prediction error associated with this reward signal (for details see [36]). We implemented a specific number of training trials, either 1, 3, 8, 15, or 30, in which dopamine-dependent plasticity was enabled at the corticostriatal synapses. After the training trials, we froze the network by turning off plasticity again and simulating an additional 50 post-training trials. This procedure allowed us to track the effects of incremental training on the network dynamics and the resulting decision behavior. Details of our CBGT model and the plasticity rule can be found in [36] and CBGT networks in the Methods section.

We next gathered the time-dependent firing rates of CBGT populations (e.g., Fig 1B) from the pre-learning and post-learning trials and analyzed them using the CLAW framework, a recently proposed computational method for tracing the instantaneous flow of neural activity through unique CBGT circuit states [32]. The method discretizes the firing rates of *N* nuclei of interest into fixed-width time bins and binarizes the mean firing rate within each bin based on whether it exceeds a predefined threshold. The network activity at each time bin is then represented as a binary *state* vector *s*_*k*_ ∈ ℝ^*N*^ (*k* = 1, 2, ···, 2^*N*^), where each element of *s*_*k*_ is either 0 or 1. In this way, the firing rate time series are converted into sequences of discrete states, each representing the pattern of CBGT activity in a time window. By aggregating transitions between states across trials, we can construct a high-probability state transition chain, i.e., the CLAW diagram, that captures the dominant trajectories of network activity throughout the decision-making process. See [32] and CLAW in the Methods section for full details of the CLAW approach.

The CLAW diagram, in the absence of learning, is depicted in Fig 2A. The accompanying table in Fig 2B provides details about the activity associated with each CLAW state. It includes the binarized firing rates of the nuclei of interest. It also reports the activation probabilities of the subthalamic nucleus (STN) and GPeA. For decisions in which a given state arises, the table reports the mean decision time (DT) and the probability that the left option was chosen. To categorize key classes of decision trajectories, the network states were partitioned into six functional *zones* (shaded blocks in Fig 2A), with the zone details listed in Fig 2C. Specifically, zone I contains the pre-stimulated state, where nuclei activity reflects baseline dynamics prior to stimulus onset. This zone functions as a *launching* phase of a decision, representing the common origin of all trials and marking the initiation of the process. Zone II comprises two loops of states that could return the trajectory to the launching zone through an initial *deliberation* process. This inner CLAW zone is critical for early evaluation of evidence, allowing the system to explore competing options rather than immediately converging on a specific action. Zones III and IV, in contrast, reflects the left and right arms of the outer CLAW, respectively, corresponding to a *committed* phase: a trial was generally locked into a specific decision once entering either arm. Exceptions to this commitment effect involves passage into zone VI, comprising a small set of states that could be reached, albeit infrequently, from either committed zone. Transitions into this zone suggests a possible *reversal* of an initial commitment, as evidenced by the decrease in the probability of left choice from zone III to zone VI (see “Prob(L)” in Fig 2B-C). Lastly, occurring after initial deliberation in zone II, zone V capturs a second-tier deliberative process. It contains a three-state loop that corresponds to significantly prolonged decision times and could lead to either choice direction. Taken together, the CLAW reveals a structured progression through three functional phases of a decision (launching, deliberation, and commitment) with different decision trials involving different paths through some or all of these phases.

**Fig 2.**
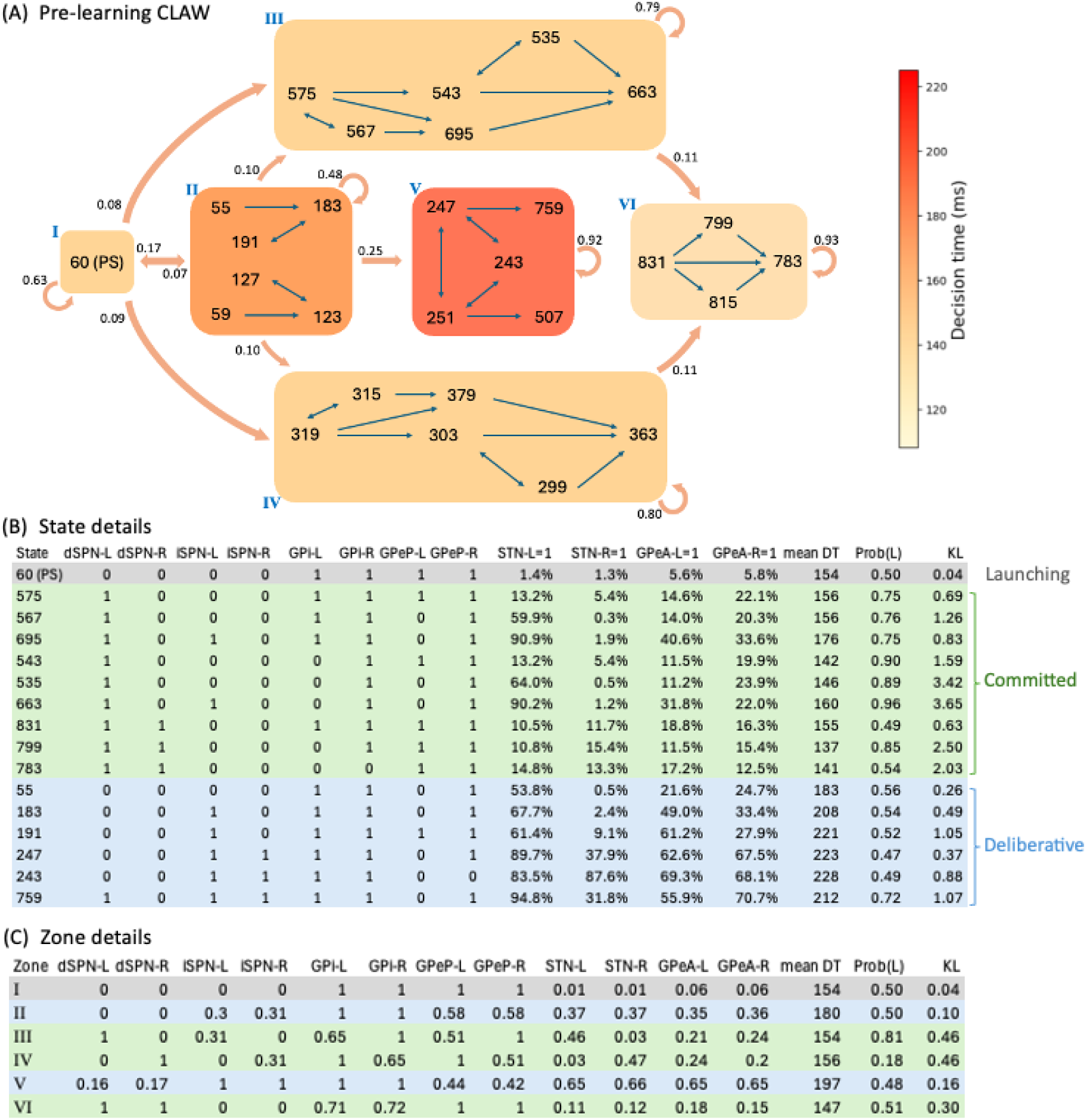
CLAW (Circuit Logic Assessed via Walks) diagram for pre-learning CBGT circuit dynamics. (A) CLAW diagram. Numbers in blocks indicate the network states, which are grouped into zones I–VI (shaded blocks). The shading of each zone indicates the mean decision time estimated over trials that visited this zone (color bar). The transition probability from one zone to another is indicated by the number near the arrow pointing from the source zone. The loop arrows represent the probability of staying in a zone and reaching the decision threshold from that zone. (B) Details of most commonly observed states. From left to right, after state labels: binarized firing rates of dSPN, iSPN, GPi, and GPeP for left (-L) and right (-R) channels; probability of activation (binarized firing rate = 1) for STN and GPeA of L and R channels; mean decision time (DT) over trials that visited each state; probability of choosing left for trials that visited each state; Kullback–Leibler (KL) divergence between the DT distributions for left- and right-choice trials that visited each state. The grey row corresponds to the initial state, or pre-stimulated state, that occurs early in each trial and never leads directly to a decision (launching), and the green and blue rows correspond to outer CLAW (commitment) and inner CLAW (deliberation) states, respectively. States in the lower half of the CLAW are not shown; these are symmetric—up to the swap of certain L and R channel binary values—with the states that lie in corresponding positions in the upper half (see the Supporting Information S1 Table for a full set of state properties). (C) Zone details. Column descriptions and row colors as in (B).

The CLAW diagram provides a compact way to distinguish two decision states—deliberation and commitment—that emerge from different patterns of engagement across CBGT pathways. Along the outer CLAW (zones III and IV in Fig 2), we observed that decision trajectories entered a committed state when activity in one action channel’s *direct* pathway became dominant due to early activation of dSPNs in the striatum (zone III for the left channel and zone IV for the right channel). This rise in dSPN activity facilitated downstream disinhibition of the thalamus, effectively locking the trajectory into commitment. Meanwhile, the indirect- and pallidostriatal-related populations, including iSPNs and both prototypical and arkypallidal cells of GPe, tended to engage more weakly or later in time, indicating that the direct pathway was the primary driver of the committed phase. In contrast, along the inner CLAW (zones II and V), decision trajectories spent more time in a deliberative regime characterized by stronger and earlier engagement of *indirect* and *pallidostriatal* pathways. In particular, the indirect pathway activity rose earlier via iSPNs (see state 183 in zone II), effectively inhibiting GPeP neurons and releasing STN from inhibition. At the same time, these changes promoted the buildup of GPeA activity, especially in the deeper deliberation stage (zone V). Here, the combined influence of iSPN and GPeA curbed the direct pathway activity, thereby keeping the system in a state where commitment was postponed while evidence continued to accumulate. The behavioral distinction between deliberative versus committed zones is captured directly in the CLAW visualization: as zone shading reflects mean decision time for trials visiting each zone, the contrast between inner zones (longer decision times) versus outer zones (shorter decision times) confirms our observation that the direct pathway drives rapid commitment while the indirect and pallidostriatal pathways shape sustained deliberation. More neural mechanistic details underlying commitment and deliberation are described in our past work [32].

Based on the sequence of phase transitions, we categorized decision trajectories into four classes (scenarios): (A) deliberative, (B) deliberative → committed, (C) committed, and (D) committed → reversal. Specifically, class A includes trajectories that exclusively engaged in the deliberative process, without transitioning into the committed phase. These trajectories followed either a short deliberative path from zone I to zone II (denoted as I → II) or a longer path that additionally visited the deeper deliberation zone (I → II → V). Importantly, despite the absence of a transition into the defined committed phase, these trajectories still displayed brief “commitment” and resulted in a choice, especially at states 759 and 507 where the dSPNs in a specific channel eventually reached a level sufficient to drive a decision outcome (see state details in Fig 2B). In class B, trajectories emerged from the initial deliberation but quickly shifted to commitment, without the need for deeper deliberation, corresponding to the paths I → II → III and I → II → IV. Other trajectories bypassed both deliberative zones and entered a commitment arm directly, i.e., I → III and I → IV, and these comprised class C. Lastly, class D captures a small number of trajectories that started in a committed zone but finally switched into the reversal zone, i.e., I → III → VI and I → IV → VI, indicating a change in the evolving dominance between action channels after an initial commitment tendency. Together, these four decision scenarios allow us to track how distinct neural temporal dynamics evolve with learning, as corticostriatal plasticity sets off downstream effects on CBGT circuit activity. In the following sections, we provide a detailed analysis of these plasticity-induced changes and their implications for decision policies.

### Learning reconfigures engagement of CBGT subnetworks

We next examined the behavioral consequences that emerge from learning, including shifts in the probability of different trajectory types and associated changes in decision time and accuracy. First, we implemented our CLAW method to analyze the post-learning trials at each plasticity stage, so as to track changes in the temporal dynamics of decision trajectories as learning progressed. Figure 3A depicts how the CLAW evolved from pre-learning (subpanel i) through early (subpanel ii) and late (subpanel iii) learning stages, eventually reaching a new, post-learning state (subpanel iv). Transition probabilities between zones with substantial changes (greater than 10% relative to the previous learning stage) are highlighted in green for increases and red for decreases. Changes in shading for each zone indicate how learning altered the average decision time associated with trajectories that entered the zone.

**Fig 3.**
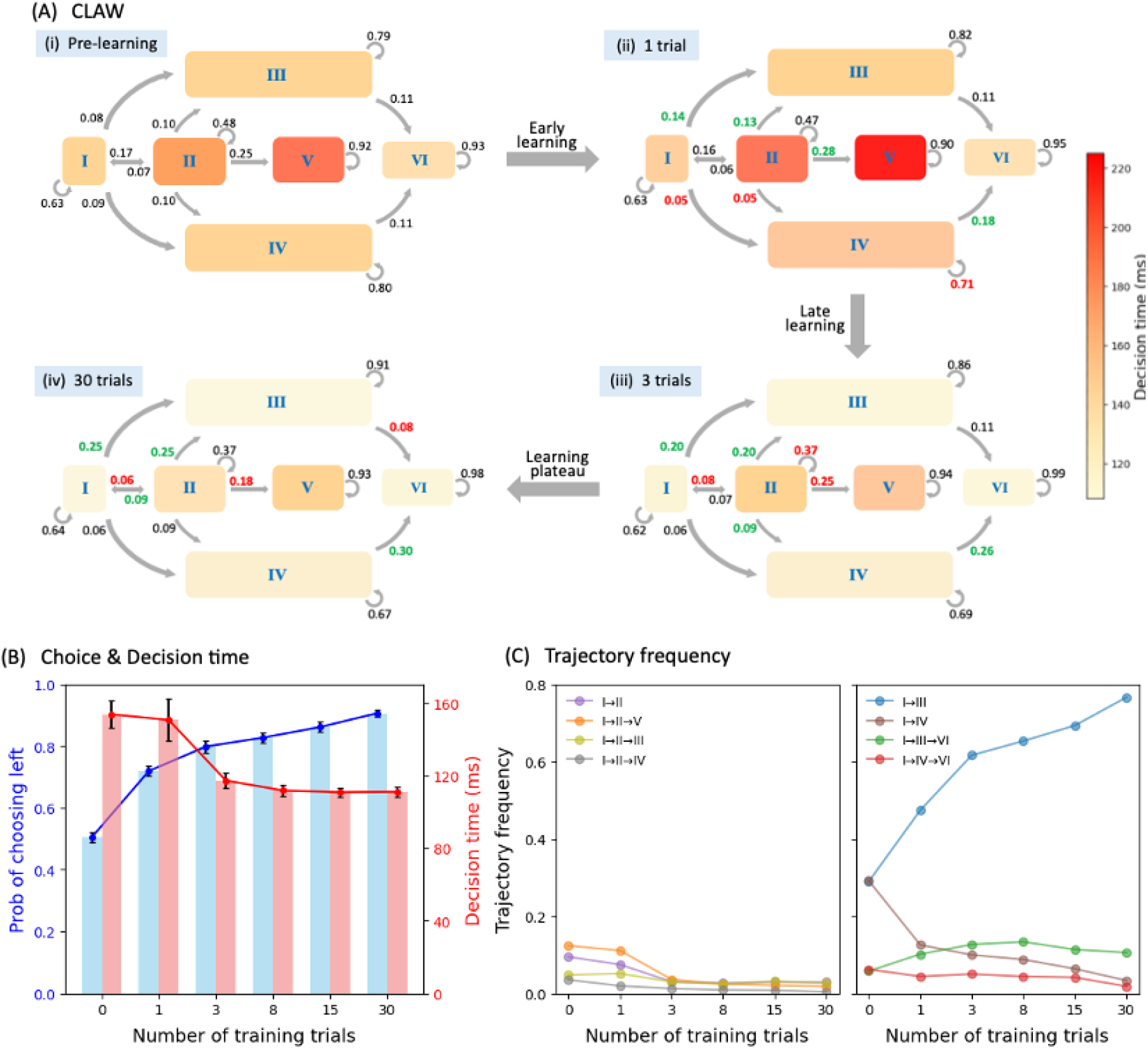
Plasticity-induced changes in decision outcomes. (A) CLAW dynamics across learning stages, from (i) pre-learning to (ii) early learning, (iii) late learning, and to (iv) learning plateau. Significant changes in the zone transition probabilities relative to the previous learning stage (greater than 10%) are highlighted in green for increases and red for decreases. The shading of each zone indicates the mean decision time over trials that visited that zone. (B) Changes in choice accuracy (blue) and decision time (red) as learning progressed. Red (resp., blue) bars indicate the mean decision time (resp., probability of choosing left) across post-training trials, with error bars representing the 95% confidence interval. (C) Changes in the relative frequency of decision types as learning progressed. Deliberative trajectories are shown in the left subpanel and committed trajectories in the right.

From pre-learning to initial early learning there was, as expected, a significant shift from launching toward more left-committed paths (I → III) and fewer right-committed paths (I → IV). For the deliberative → committed trajectories, the initial deliberation led to more transitions to left-committed states (II → III) and fewer transitions to right-committed states (II → IV). The reversal path also showed an upward trend, with more initially right-committed trajectories switching toward the left (IV → VI). These shifts suggest that even after single-trial experience, the system began to favor the rewarded direction. However, an exception occurred in the purely deliberative trajectories, where the entry into the initial deliberation stayed almost unchanged (I → II) and the transitions to the second deliberation increased (II → V). Darker shading of zones II and V further indicates an extension of the deliberation time. These changes suggest that, despite the increasing tendency to rapidly make left decisions, the deliberative types of decisions preserve a degree of caution after limited learning, requiring more thorough and extended evaluation to finalize their decisions.

Comparing the CLAW after three training trials to that after one training trial, several notable shifts emerged that aligned with the system’s progression from early to late learning. The most significant shift was the 50% drop in the probability of entering the first deliberation zone (I → II), indicating increased confidence that allowed the system to avoid prolonged uncertainty. Moreover, we observed a decline in the probability of lingering in zone II (loop arrow), giving way to more direct flow into committed zones. In addition, the likelihood of proceeding to a second deliberation (II → V) also diminished, and the time spent in both deliberation zones significantly decreased (zone shading). All these changes suggest that the deliberative phase, previously more prominent during early learning, was disfavored and compressed after sufficient experience, shifting the system toward a more streamlined mode of decision-making.

After three training trials the pace of learning began to decelerate (8- and 15-trial CLAWs not shown). Changes observed in the previous trials, such as reduced deliberation and increased commitment, were slowly refined, but no qualitatively new changes emerged. With 30 training trials the trends towards direct commitment and towards correction of initially right-oriented trajectories, alongside reduced initial and second deliberation, all continued and stabilized. Overall, after initial rapid improvements, the system largely saturated its capacity for adaptation, and further training produced limited returns. Together, the CLAW series in Fig 3A reveals that learning progressively biases the CBGT circuit dynamics toward faster, reward-directed decisions.

Further supporting this progression, Fig 3B-C illustrates how the overall decision time, choice accuracy, and trajectory type evolved over the course of learning. Choice accuracy (the probability of choosing the rewarded left option; blue symbols in panel B) climbed steadily from chance (50%) before learning to 90% after 30 training trials. Concurrently, decision time (red symbols) decreased substantially with learning. Early in learning, mean decision time remained high and variable due to prolonged deliberation (consistent with the inner CLAW shading from subpanels i to ii in Fig 3A); however, with further training, decision times significantly decreased with reduced variance and rapidly stabilized. In Fig 3C, we quantified the extent to which training gradually limited deliberative decisions and favored left-committed ones. The relative frequencies of deliberative trajectories plateaued early, whereas those of committed trajectories continued to exhibit small refinements. This pattern is also consistent with shifts observed in zone transitions across the CLAW in Fig 3A. Overall, these results illustrate that dopamine-driven plasticity at the corticostriatal synapses efficiently restructures the CBGT dynamics manifested as trajectories progressing through the CLAW, such that the system spent less time in indecisive deliberation and more often moved straight toward the rewarded choice, yielding faster and more accurate decisions.

To understand the specific CBGT subnetworks that were responsible for these learning-induced changes, we next analyzed the engagement of *control ensembles* identified within CBGT circuits. We have previously shown that, in the absence of learning, there exist three low-dimensional subnetworks, referred to as control ensembles. Each control ensemble is associated with specific configurations of the decision policy parameters in the drift-diffusion model [29, 31, 32]. To characterize these ensembles, two aspects of CBGT activity were considered: (1) the overall firing rates across the left and right action channels (i.e., sum of the firing rates in each cell population across both channels, denoted as L+R), and (2) the bias in activity towards one action (i.e., difference in the firing rates of each population between left and right channels, denoted as L–R). Using canonical correlation analysis (CCA), we captured three components—*choice, responsiveness*, and *pliancy* —that maximally correlated variation in the space of CBGT activity and variation in the space of DDM parameters. Here we reproduced these control ensembles as described in [32], with the aim of analyzing how the dopaminergic learning process modulated their engagement, thereby shaping the resulting behavioral outcomes. Prior work [29, 31, 32] and Control ensembles in the Methods section provide more methodological details of this approach.

We visualize the ensembles as they were manifested in the CBGT circuit and the DDM in Fig 4. The choice ensemble (panel A) is driven by the between-channel activity differences across all populations. This subnetwork encodes the dominance of one action channel over the other and primarily influences the drift rate *v*. Color coding indicates the direction of influence, with green for cell types where larger differences between channels align with increases in left-directed drift rate (i.e., larger *v*), indicating more rapid evidence accumulation towards the rewarded option, and pink for the opposite. We do not show the loadings of overall activity across channels for the choice ensemble because they are extremely weak, meaning that overall activity has very little impact on the choice-related parameters of the DDM process. In contrast to the choice ensemble, responsiveness (panel B) and pliancy (panel C) represent subnetworks that are primarily defined by the overall CBGT activity across both channels, with minimal loading on differential activity (not shown). The responsiveness ensemble features strongest loading on the corticothalamic and direct pathways. It modulates both how quickly evidence evaluation begins (onset time *t*) and the level of evidence needed for a commitment to a decision (boundary height *a*), and hence the overall response speed. The strongest loadings in the pliancy ensemble (panel C) arise in the overall activity in components of both the indirect and pallidostriatal pathways. As with responsiveness, pliancy is strongly associated with *t* and *a*, but with an opposite direction of effect on *a*, reflecting how effectively the accumulated evidence is translated into decision execution.

**Fig 4.**
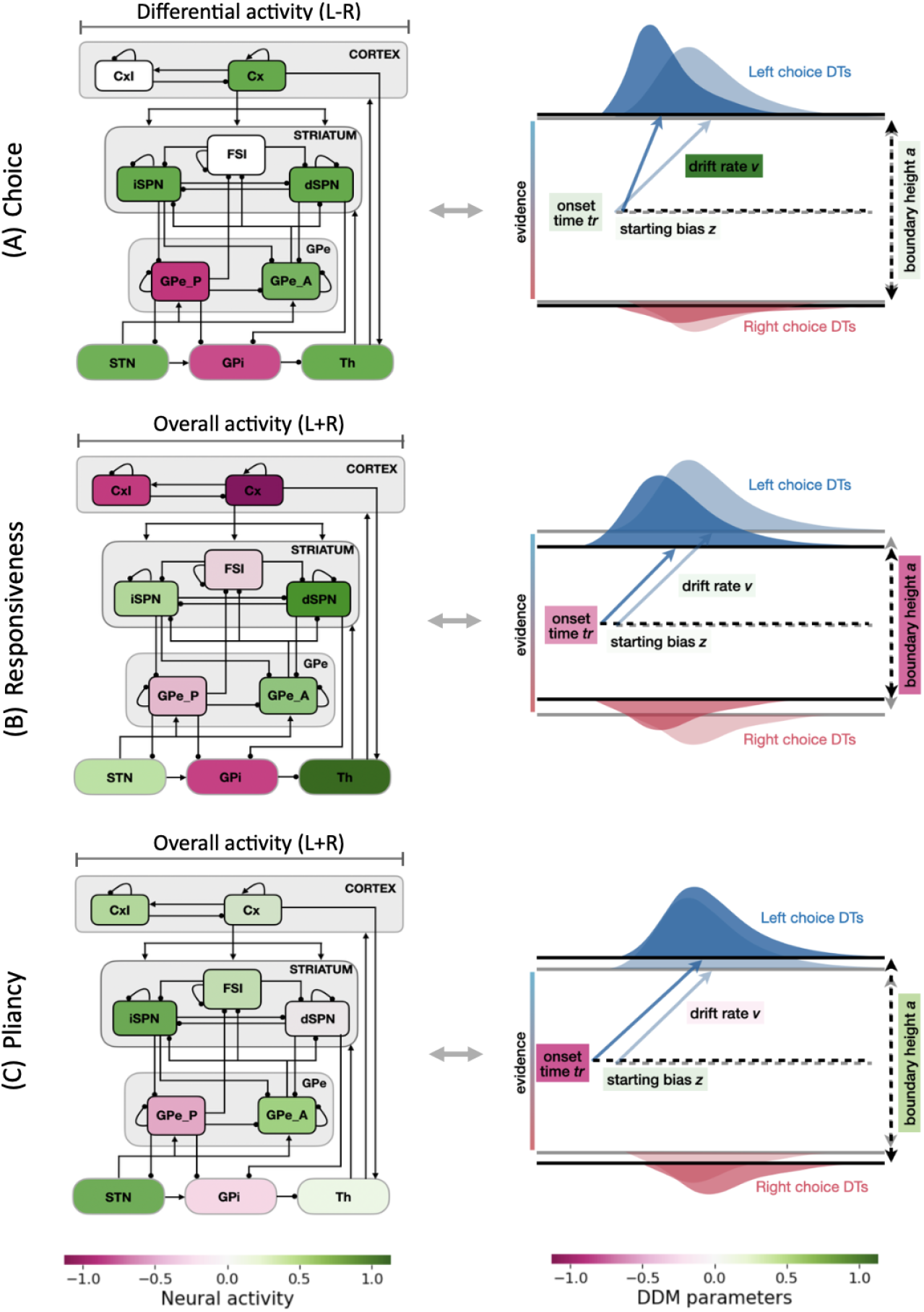
Control ensembles and DDM correlates identified by canonical correlation analysis (CCA). Panels A, B, and C represent the relation between the firing rate of each population in the CBGT circuit and the DDM parameters obtained in the first three canonical components, labeled as choice, responsiveness, and pliancy, respectively. Each left subpanel represents neural contributions: for the choice ensemble (panel A), this is the differential activity between the left and right channels (L–R); for the responsiveness and pliancy ensembles (panels B and C), this is the overall activity summed across both channels (L+R). The other aspect of activity for each ensemble (L+R for choice and L–R for responsiveness/pliancy) is not shown, as it exhibits minimal impact on DDM parameters. Each right subpanel depicts the associated changes in the DDM parameters, as well as the resulting effect on the DT distributions. In all panels, green shading denotes increases and pink denotes decreases.

Since the three control ensembles approximately decompose the complex CBGT circuit into specific subnetworks, i.e., differential activity in all pathways (choice), overall activity in corticothalamic and direct pathways (responsiveness), and overall activity in indirect and pallidostriatal pathways (pliancy), we next investigated changes in subnetwork engagement across incremental training. This analysis was meant to help explain how feedback-driven plasticity progressively reconfigured the within-trial pattern of engagement of these CBGT subnetworks. To this end, we translated the time series of population firing rates into the time course of individual control ensembles (see Control ensembles in the Methods section for details). Given that decision times varied across trials, we performed averaging by aligning trials separately to both cue onset and decision time to capture both the early launching and late committed dynamics critical for the decision process, as shown in Fig 5. We focused on two temporal windows: the first 50 ms after cue onset, corresponding to the launching phase, and the the final 30 ms prior to decision, corresponding to the committed phase. The intermediate period, consisting of the remaining launching phase and/or deliberative phase, was not considered due to high variability in its duration, although we inferred changes evident at the end of deliberation from values at the onset of the committed phase. Figure 5 shows the control ensemble engagement averaged across all trials (A panels), as well as separately for left-choice (B panels) and right-choice (C panels) trials. Corresponding ensemble time courses broken down by decision-trajectory types are provided in the Supporting Information S3 Figure.

**Fig 5.**
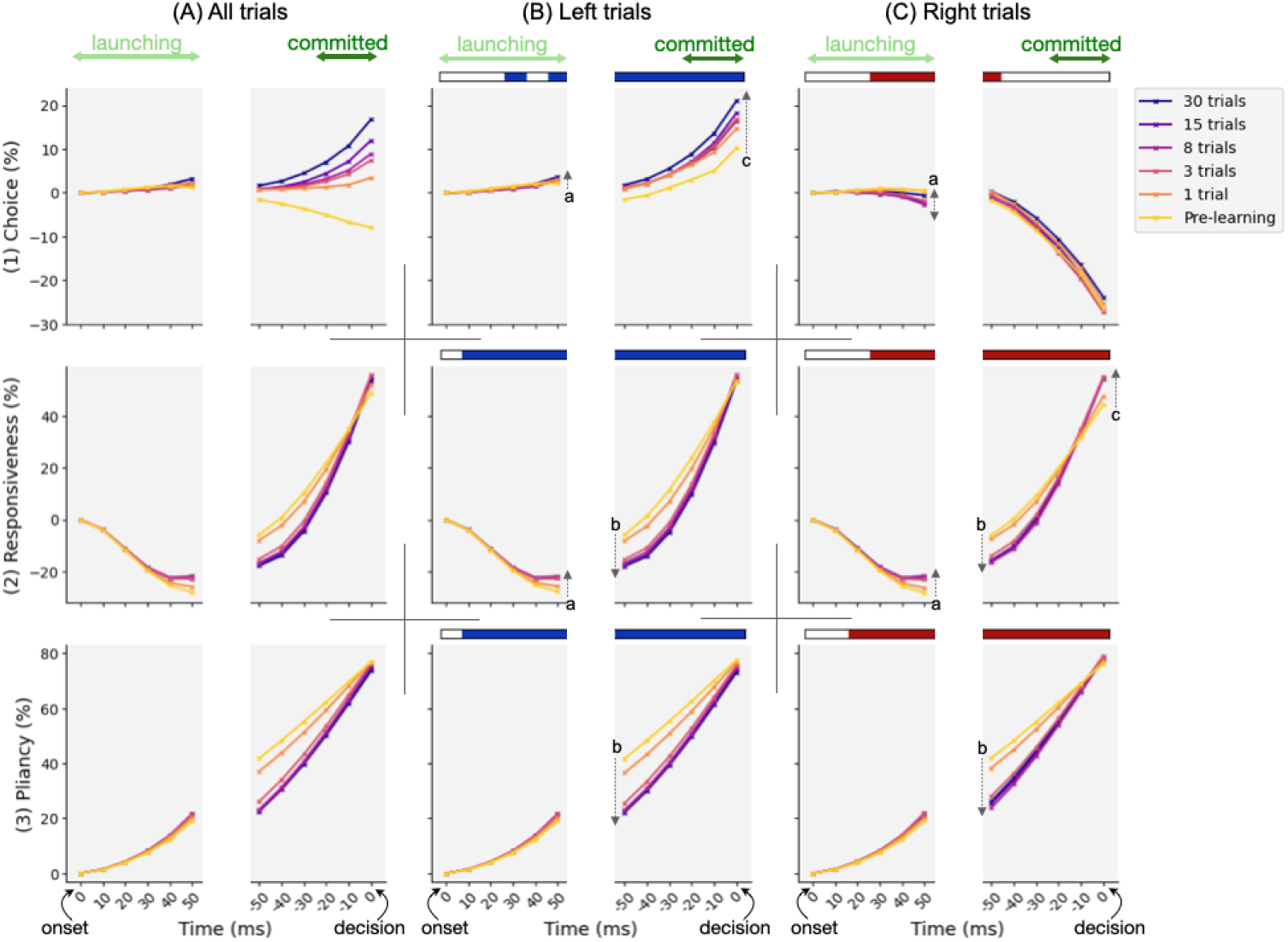
Time evolution of control ensemble engagement at different learning stages. Trace hue represents learning stage, from pre-learning (yellow) to 30 trials of training (purple). Plotted values are percentage changes in engagement relative to cue onset, averaged across (A) all trials, (B) left-choice trials, and (C) right-choice trials. Each panel includes traces aligned to cue onset (time 0 on the left end of the x-axis) as well as traces aligned to decision time (time 0 on the right end). The first 50 ms after cue onset corresponds to the launching phase, and the final 30 ms prior to decisions corresponds to the committed phase. The intermediate period, comprising the remaining launching phase and/or deliberative phase, was omitted due to variability in its duration across trials, although we inferred changes evident at the end of deliberation from values at the onset of the committed phase. Shaded horizontal bars above each panel of columns B and C indicate time bins in which the learning effect on the corresponding control ensemble was statistically significant. Dotted arrows in each panel mark the direction and magnitude of the learning effect during launching (arrows a), deliberation (arrows b), and committed (arrows c) phases.

The time course of choice ensemble engagement shows a modest deviation from baseline during the launching period, becoming slightly positive for left decisions (panel B1) and negative for right decisions (panel C1). This early bias blossomed into a strong directional commitment that increased throughout the committed phase. Here we note that during commitment, the pre-learning choice engagement tended to decline (yellow trace in A1). This pattern arose from chance trial-to-trial fluctuations in the inputs that, when averaged, produced this trend. Nevertheless, it was immediately corrected after a single training trial. For the responsiveness ensemble engagement (row 2), it initially decreased following cue onset, reflecting an early suppression in the corticothalamic and direct pathways. This suppression was followed by a rapid increase up to the decision time, consistent with the enhanced engagement of this subnetwork when approaching to a decision. Pliancy ensemble engagement, by contrast, steadily increased throughout the course of decisions (row 3), reflecting a sustained influence of indirect and pallidostriatal pathways. Given the opposing influence of responsiveness and pliancy on boundary height (Fig 4B-C), their divergence during launching implies complementary control that supported more deliberation, whereas their concurrent rise near commitment indicates competing influences as the process moved toward action selection, which will be discussed in more detail later.

As we looked at traces across training (trace color from yellow = pre-learning to purple = 30-trial training), several phase-specific effects of learning are evident. To identify when learning exerted a statistically significant effect on control ensemble engagement, we fit a mixed-effects regression model. The model included fixed effects for learning stage and trajectory type over time, along with a random effect of network variability (see Learning effect in the Methods section). Time bins with a significant learning effect are indicated by shaded horizontal bars above each panel of left-trial and right-trial groups, and dotted arrows within the panel illustrate the direction and magnitude of the effect during launching (arrows a), deliberative (arrows b), and committed (arrows c) phases. Overall, we observed that both 1 and 3 trials of training resulted in clear shifts in the dynamics of the three control ensembles. After 3 trials, the traces of both responsiveness (row 2) and pliancy (row 3) began to converge and they showed little further change with continued training, suggesting that the underlying global network dynamics, and its contribution to decision time, quickly reached a relatively steady state. In contrast, the choice ensemble (row 1) continued to show changes beyond 3 trials of training, implying that competition between action channels, potentially contributing to decision accuracy, required extended experience to stabilize. Such modulation of control ensemble engagement matches our early/late/plateau learning stages, and is consistent with the behavioral trends shown in Fig 3.

Examining phase-specific effects, during the 50 ms launching phase after cue onset, learning amplified the initial subtle bias: in left-choice trials, the choice ensemble slightly increased in the rewarded direction, whereas in right-choice trials the early rightward bias was non-monotonic across learning stages (arrows a in B1 and C1). Additionally, during this launching period learning raised the negative responsiveness curve (arrows a in B2 and C2), while the pliancy ensemble was almost unaffected. These shifts at the beginning of the decision process suggest that learning sharpened initial directional bias while concurrently adjusting global network drive, primarily by shifting relative neuronal engagement away from cortex and towards the direct pathway (specifically, dSPNs and thalamus) via an increase in responsiveness, which prepared the system for faster engagement without immediately triggering commitment.

During the subsequent deliberation window, however, learning led to a transient suppression of responsiveness (arrows b in B2 and C2) rather than a continuation of its initial increase. This suppression acted as a regulatory mechanism: the temporary constraint on responsiveness would limit premature commitment and support more sustained deliberation. A similar suppression was observed for pliancy (arrows b in B3 and C3), which was associated with the indirect and pallidostriatal pathways. Although this reduction may seem counterintuitive given their role in deliberation, it likely reflects a coordinated downscaling of global network drive during this phase, allowing relative differences between action channels (i.e., choice ensemble) to continue accumulating while overall activity was prevented from ramping to levels that would trigger early commitment. Consistent with this pattern, combined influence of responsiveness and pliancy on boundary height reveals a net increase in the decision threshold during this window, effectively preserving deliberation (discussed in the next subsection).

In the final 30 ms committed phase before a decision, learning produced a pronounced increase in choice ensemble engagement for left trials (arrow c in B1) and induced a rebound of both responsiveness and pliancy to, or slightly above, their pre-learning levels at the moment of decision (B2, B3, C2, and C3). This late-phase learning modulation marked a transition from temporary restraint to decisive engagement, in which restored global drive, together with an amplified choice signal, facilitated rapid and reward-directed action commitment.

Taken together, all the three control ensembles showed significant modulation with learning. On the one hand, the growing early choice bias pushed the system toward the rewarded channel faster. On the other hand, the temporary mid-phase suppression of overall activity, especially via thalamic and dSPN populations, enforced caution and counteracted the increasing between-channel asymmetry. Finally, in the commitment phase, all subnetworks worked together to trigger a rapid decision once sufficient evidence had accumulated. Thus, these dynamics indicate that learning implemented a coordinated tuning of both global (network-wide) and selective (between-channel) activity within CBGT circuits so as to manage a balance between caution and commitment. To derive a more quantitative assessment of the impact of learning, we next analyzed more directly how the plasticity-driven tuning of CBGT subnetworks translated into changes in decision policies.

### CBGT subnetwork interplay reshapes decision policy

For our final analysis, we translated the learning-induced reconfiguration of CBGT subnetworks into an algorithmic description by projecting to the corresponding modulation of key parameters of the drift-diffusion model that describe the properties of the decision policy of the CBGT network. We used our CCA results (Fig 4) to map the time series of control ensemble engagement (Fig 5) to the DDM space, and examined the within-trial evolution of drift rate and boundary height at each learning stage (see Control ensembles in the Methods section). As shown in Fig 6, changes in drift rate (row 1) were highly correlated with the choice ensemble dynamics (row 1 in Fig 5), as drift rate was primarily driven by the choice ensemble (Fig 4A). Hence, as learning advanced, evidence accumulation became increasingly biased and, in the commitment phase, accelerated more rapidly toward the rewarded option.

**Fig 6.**
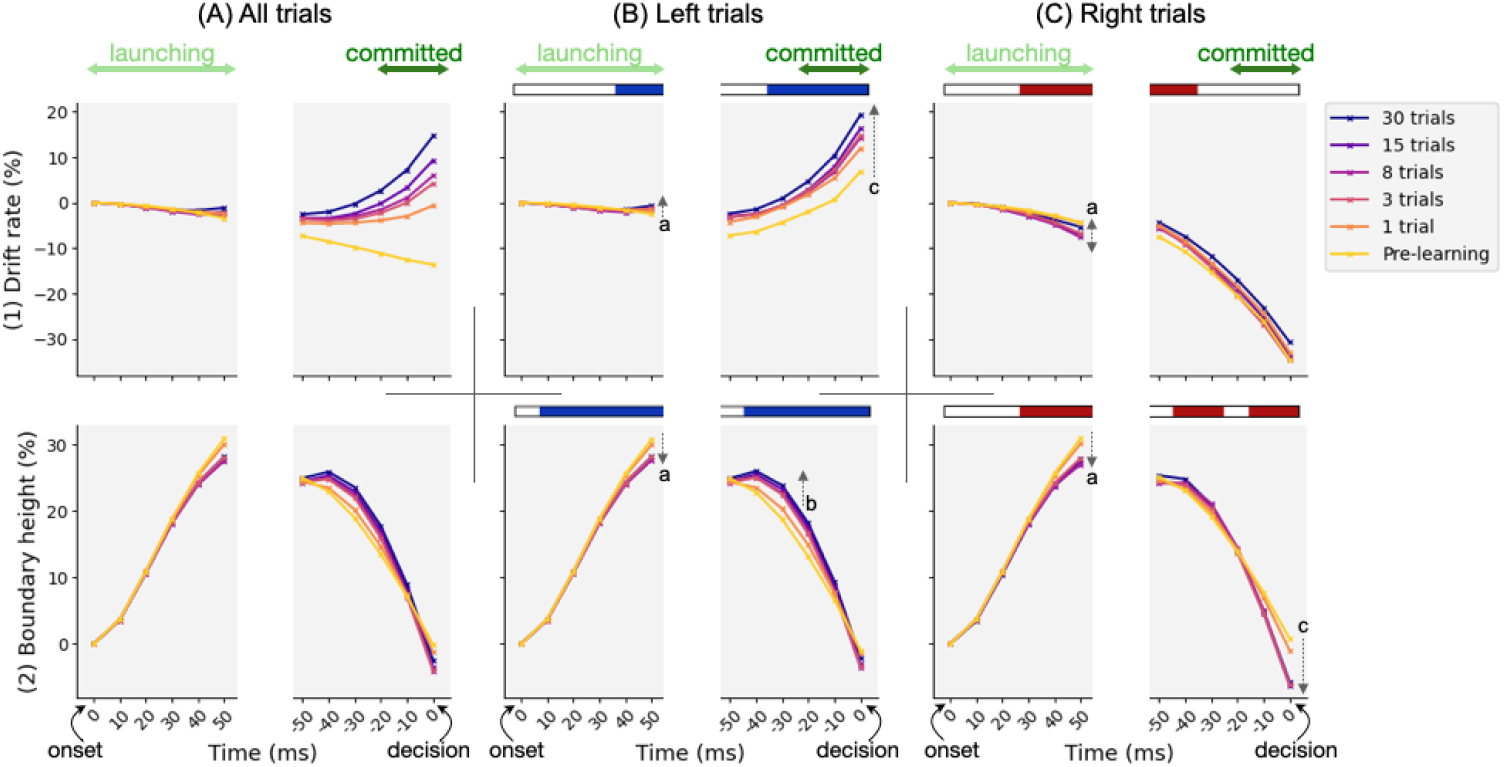
Time evolution of drift rate and boundary height at different learning stages. Traces were aligned to cue onset (time 0 on the left end of the x-axis) and decision time (time 0 on the right). Shaded horizontal bars above each panel of columns B and C indicate time bins in which the learning effect on the corresponding DDM parameter was statistically significant. Colors, arrows, and notations as in Fig 5.

Unlike drift rate, changes in boundary height (row 2) resulted from a more complex interplay between responsiveness and pliancy ensembles, because increased responsiveness tended to lower the threshold, whereas increased pliancy tended to elevate it (Fig 4B-C). To disentangle these effects, we investigated their relative contributions to decision boundaries (Fig 7), computed by multiplying each ensemble’s engagement by its loading on boundary height and normalizing the two terms to obtain proportional contributions. The plotted curve represents the proportion of responsiveness contribution; the complementary proportion (1 − responsiveness) corresponds to pliancy. Combining this decomposition with boundary height traces in Fig 6, we observed that early in the decision, responsiveness (which decreased; see row 2 in Fig 5) accounted for a larger share of the initial rise in boundary height, consistent with the initial decrease in corticothalamic and direct pathway activity that also raised the onset time to trigger the evidence accumulation process after cue onset. As the decision unfolded, the relative contribution of responsiveness decreased while pliancy became more dominant (Fig 7), shifting control toward indirect and pallidostriatal influences that maintained the high threshold during deliberation (row 2 in Fig 6). Near commitment, responsiveness regained its influence (Fig 7), driving boundary collapse (row 2 in Fig 6) and enabling action selection.

**Fig 7.**
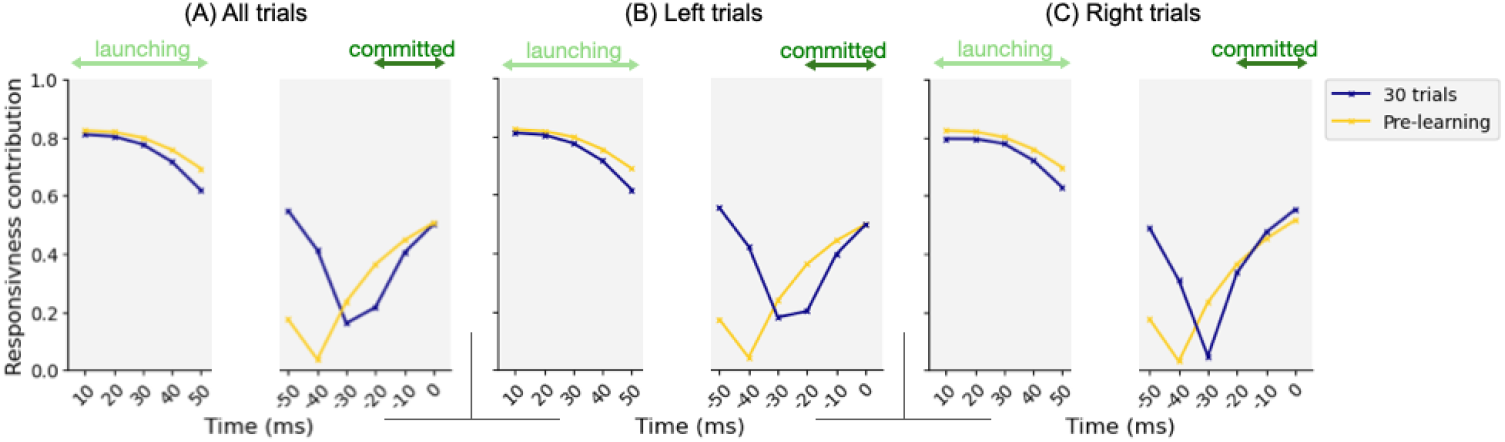
Contributions of responsiveness and pliancy to boundary height. Traces were aligned to cue onset (time 0 on the left end of the x-axis) and decision time (time 0 on the right). At each time bin, the contribution of each ensemble to boundary height was computed as ensemble engagement (Fig 5) multiplied by its loading on boundary height *a* (Fig 4B-C), and the two terms were normalized to yield proportional contributions. The plotted traces show the responsiveness proportion; the complementary proportion, (1 − responsiveness), corresponds to pliancy.

Further, comparing post-learning (blue) with pre-learning (yellow) traces, we found that learning dramatically curtailed the drop in the responsiveness contribution during the intermediate period of the decision process leading up to commitment, consistent with the mid-phase suppression of responsiveness engagement (arrow b in Fig 5B, row 2). This suppressed responsiveness explains the learning-induced increase in boundary height during deliberation (arrows b in Fig 6, row 2), helping to preserve caution as drift increased (row 1) and thus reducing premature commitment. Hence, these results reveal that over the course of a decision, different CBGT subnetworks dominate at different phases and trade their engagement over time. This phase-specific control regulates the non-monotonic pattern of boundary height changes, thereby allowing for the possibility of deliberation while also promoting timely commitment.

To summarize our results so far, we developed a dynamic DDM fitting approach that captures how decision policy parameters change as individual decisions progress (see also [32]). Unlike static DDMs that assume a time-homogeneous process with fixed parameters throughout the decision, our method allows drift rate and boundary height to take different constant values within different decision phases, with values specific to each trajectory type (as identified by CLAW in Fig 2), allowing the evidence accumulation process to reflect the dynamic nature of neural activity over the course of a decision. Full description and validation of this approach are provided in Dynamic DDM fitting of the Methods section. It remains unclear how this dynamic process is influenced over the course of learning. Figure 8 shows each decision trajectory projected to the joint drift rate and boundary height space, i.e., (| *v* |, *a*), across pre-learning (light hue), early learning (medium hue), and late learning (dark hue) stages. Each square along a trajectory represents the parameter values at the corresponding CLAW zone, and background manifolds show the mean decision time predicted by DDM simulations for each (| *v* |, *a*) combination.

**Fig 8.**
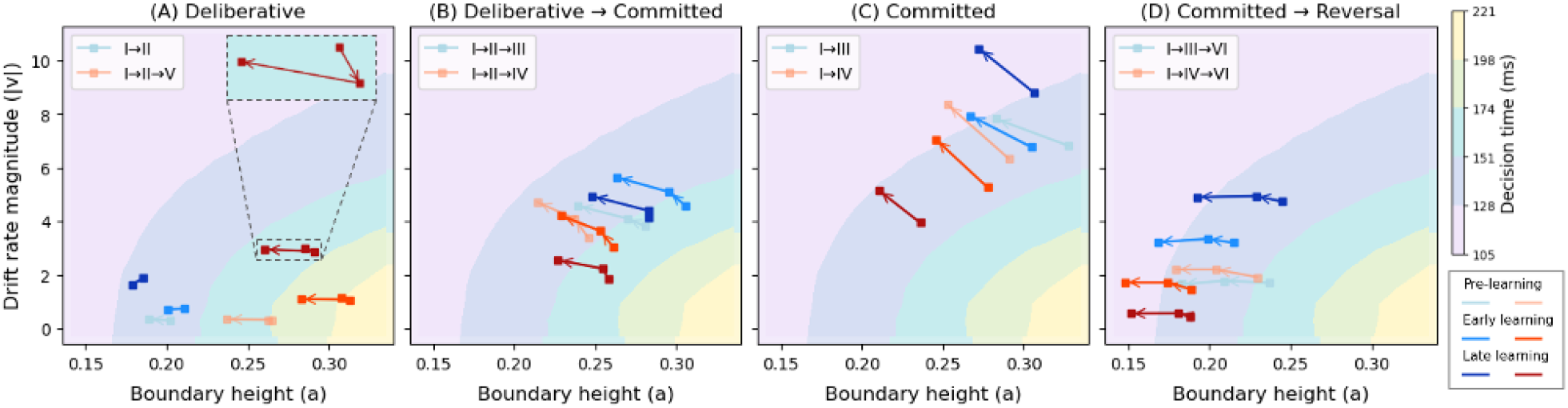
Plasticity-induced changes in the decision trajectories projected to DDM space. Trajectories were mapped onto the (|*v*|, *a*)-space by a dynamic DDM fit. Within each panel, each trajectory type is represented by a color family (blue or orange). Each square along a trajectory represents the parameter values at the corresponding CLAW zone (either two or three zones per trajectory, as indicated in legends). For example, the dark blue trajectory in panel C flows upward and leftward from zone I to zone III. Line darkness represents plasticity stages: pre-learning (light), early learning (medium), and late learning (dark). Background manifolds indicate the average decision time associated with each (|*v*|, *a*) combination, estimated by simulating the DDM.

Across panels A–D of Fig 8, the four trajectory classes occupy distinct regions of the (|*v*|, *a*)-space that reflect their different strategies. In panel A (deliberative class), trajectories start with relatively small |*v*| and moderate *a*, consistent with slow, uncertain accumulation that supports potential returns to earlier zones. In panel B (deliberative → committed class), trajectories move upward and rightward relative to panel A, indicating more decisive evidence accumulation together with elevated boundary heights that still allow for initial exploration. In panel C (committed class), they shift further upward and rightward, clustering at larger | *v* | and higher *a*, reflecting fast, directed accumulation accompanied by enhanced thresholds to prevent overly rapid commitment in the face of stronger drift. Finally, trajectories in panel D (committed → reversal class) occupy a distinct region with lower | *v* | and lower *a* than purely committed trajectories, reflecting weaker momentary evidence and reduced thresholds that together facilitate switching rather than sustained commitment. Accordingly, as decision types transition from uncertain, deliberative trajectories to more certain, committed ones, the distribution of trajectories shifts from the lower-left toward the upper-right of the (| *v* |, *a*)-space. This transition indicates two coordinated changes: (i) an increase in drift rate magnitude, corresponding to faster evidence accumulation toward the rewarded option, and (ii) a concurrent increase in boundary height that compensates for the accelerated drift to avoid premature commitment.

Secondly, within each trajectory (i.e., each connected line), parameters also evolve systematically across CLAW zones. Most trajectories show a general trend of increased drift rate magnitude and decreased boundary height as they flow across zones during the decision process. Interestingly, both deliberative and reversal trajectories (panels A and D) show less of an increase in drift rate than the other classes, including some weak non-monotonicity (panel A, inset). Indeed, the lower overall drift rates in the reversal case suggest that even though trajectories’ initial zone paths match those of committed trajectories, there are in fact early warning signs of the upcoming switch. Within CBGT circuits, this distinction would correspond to more similar activity levels across action channels than in pure commitment.

Finally, zooming in within each trajectory type (i.e., the same color group in each panel), we observed distinct learning-driven modulations on the two DDM parameters. Drift rate shows a relatively monotonic increase across learning stages (becoming more positive for left trajectories such as I → III and less negative for right trajectories such as I → IV), indicating a consistent strengthening of reward-directed bias. In contrast, boundary height evolves more dynamically across stages in ways that depend on the trajectory type. For trajectory types involving deliberation, boundary height is elevated in early learning and then declines in the later stage (Fig 8A-B). This pattern corresponds to longer decision times, as reflected by the background manifolds, and also aligns with the increased time spent in the inner CLAW (see subpanels i→ii in Fig 3A). For trajectories with a more direct progression to commitment, early learning produce decreases in boundary height (Fig 8C-D), consistent with shorter decision times illustrated by the arrows in Fig 6B-C. These decreases continue later in learning (darkest lines) for right trajectories but less so for left trajectories. In these cases, limited experience is sufficient to promote a more confident, left-committed strategy, whereas deliberative strategies remain comparatively cautious.

Taken together, these results indicate that learning increases reward-directed drive (via the drift rate) while adjusting caution (via boundary height changes) in a trajectory-dependent manner, effectively tuning the balance between deliberation and commitment. In this view, boundary modulation acts as a compensatory control that either preserves caution when drift increases (deliberative decisions) or relaxes it when early evidence is already decisive (committed decisions), yielding distinct policy refinements across decision classes.

## Discussion

Adaptive decision-making depends on the brain’s ability to learn from experience and to adjust decision policies that guide how future actions are chosen. Here we examined how such flexibility can be controlled by CBGT circuits through coordinated adjustments of a set of underlying control ensembles. These subnetworks regulate specific aspects of the emergent decision policy, such as evidence accumulation rate and selection threshold, providing a mechanism by which the decision process can dynamically adapt to environmental feedback. In this work, we integrated multiple computational approaches, including a generative large-scale spiking neural network model and high-dimensional machine learning techniques, to uncover how plasticity within CBGT circuits can have a multifaceted effect on cognitive decision processes. In particular, we analyzed how the reward-driven dopamine-mediated plasticity at corticostriatal synapses induces changes in the competition between deliberation and commitment within the time course of a single decision (Fig 3). We next looked at the neural activity driving these changes, as manifested in the temporal patterns of CBGT control ensembles during individual decisions (Fig 5). At a more abstract level, we showed how these patterns could be represented in terms of the parameters of an evidence accumulation process (Fig 6) and how plasticity-induced changes in the balance among control ensembles translated into changes in aspects of decision policy (Fig 7), with different effects arising in different decision trajectory classes (Fig 8). Overall, these results show that feedback-driven changes in control ensembles produce distinct adjustments to the underlying decision policy, reshaping how repetitions of similar sensory evidence shift successive decisions to increase reward.

A particularly notable aspect of our findings was the phase-dependent effect of plasticity on decision dynamics (see Fig 5). Although responsiveness responded more rapidly to cue onset after learning, facilitated by dopamine changes in the direct pathway, its increased engagement was not sustained throughout the decision process. Instead, during deliberation, responsiveness and pliancy were both transiently suppressed relative to the pre-learning baseline. Functionally, this pattern supports a separation between the early expression of tendencies and later commitment: learning can increase sensitivity to value early on, while preserving a window of time in which incoming evidence can still shape the choice and prevent premature commitment. In this way, the decision process can move toward exploiting learned values without eliminating the possibility of exploring or evaluating incoming evidence. This interpretation aligns with prior experimental work showing that phasic dopamine does not eliminate the system’s capacity for exploration. For example, phasic optogenetic stimulation of SNc dopaminergic neurons in mice increased exploratory behavior during initial exposure to a novel environment [37], and enhanced dopamine signaling in humans boosted exploitation while preserving the capacity for random exploration [38].

The suppression of direct pathway drive during deliberation through learning did not, however, slow decisions overall. Rather, the left–right channel differences accumulated over the course of learning prompted earlier transitions to the committed phase, thereby contracting the deliberation window. Learning produced a sharper rebound of overall CBGT activity during commitment (Fig 5, rows 2-3), which synergized with a pronounced late-stage increase in left–right differential activity (row 1) to facilitate decisive selection. This coordination points to regulation of the timing of CBGT subnetwork engagement across key phases of each decision, implying that learning not only optimizes *what* choice is favored but also adjusts *when* different populations control the process. Related timing results—though they may vary across tasks and reward conditions—are broadly consistent with our interpretation. For example, optogenetic activation of direct pathway neurons in mice can initiate learned anticipatory licking [39], which would correspond to increased responsiveness and hence reduced boundary height during launching time in our model. Conversely, transient inactivation of the unilateral striatum during either the first or second half of the decision process induced a significant ipsilateral choice bias in mice [40], consistent with the increased activity asymmetry between competing action channels that drive the choice ensemble.

In light of the diverse temporal patterns of CBGT activity observed over the course of individual trials, our work here proposes a unified and testable account of how learning can reshape the entire progression of the decision-making process. One behavioral prediction is that decision times should be more variable early in learning, consistent with less stable commitment and greater exploration, and then become both faster and less variable as learning consolidates (cf. Fig 3B). Related motor learning studies similarly indicate that behavioral variability is actively regulated during learning: it is often elevated early, but decreases as reinforcement learning progresses and identifies high-reward action variants, while remaining flexibly modulated by recent outcomes and task demands [41–43]. This pattern for decision times could be tested by comparing within-subject decision time distributions (e.g., variance and quantiles) across early versus late learning blocks. At the neural level, the temporal analyses of core cell populations in CBGT subnetworks (summarized in the Supporting Information S4 Figure) predict a concurrent shift toward stronger choice-selective signals (increasing L–R separation) alongside reduced global activity (decreasing L+R) during deliberation, consistent with the idea of maintaining caution despite an increased value-driven preference. Together, these predictions link learning-related changes in behavioral outcomes to experimentally measurable changes in population dynamics across the basal ganglia circuitry.

The CBGT mechanisms revealed by our analysis also support the emerging perspective that striatal direct and indirect pathways act cooperatively to regulate the process of value-based decision-making [5, 13, 44–48]. Note that this idea does not contradict the classic view of their opposing contributions [11, 12, 15], but rather highlights that opposing roles do not imply anti-correlated activity. We add to this point by proposing a more nuanced competition and coordination across the time course of a decision and by suggesting that differences in their interactions may map to distinct tradeoffs of deliberation versus commitment. Specifically, in our simulations, the direct pathway became increasingly active in initiating responses and driving commitment as learning advanced, whereas the indirect pathway gained strength in intermediate phases to maintain some capacity for deliberation (cf. Fig 7). Moreover, note that the non-dominant component in each phase was not idle, but played a regulatory role in shaping decision dynamics. During initiation, although it remained subthreshold, iSPN activity started to ramp up (consistent with [44]), facilitating activation of the overall network necessary for launching the evidence accumulation process. During deliberation, dSPNs maintained some activity that mitigated the indirect pathway’s dominance on deliberation (extending the balance mechanism raised in [46]), although this contribution became less important with learning as the duration of deliberation diminished. During commitment, the indirect pathway maintained a modest yet stabilizing role (supported by [47] where stimulation of iSPNs altered selection timing without impairing the ability to execute selection). This sequence of direct-indirect interplay highlights that both pathways contribute to each decision phase in the face of learning-driven changes in internal states, offering a potential reconciliation between traditional frameworks of their antagonistic roles and emerging findings of their cooperative dynamics.

The “upward mapping approach” has now been used repeatedly to fit CBGT-driven decision outcomes to a normative DDM model, leading to the consistent identification of three CBGT control ensembles [29, 31, 32]. Based on this control ensemble analysis, we translated the time series of CBGT firing rates into the temporal evolution of individual DDM parameters (Fig 5) and projected these dynamics into trajectories within the parameter space defined by drift rate and boundary height (Fig 6). This mapping allowed us to interpret circuit-level dynamics in terms of their algorithmic roles in evidence accumulation. In particular, we observed that dopaminergic plasticity within the CBGT circuit induced distinct effects on drift rate and boundary height. Drift rate, which was tightly correlated with choice, became increasingly rapid toward the rewarded direction both over time within a trial (“horizontally”) and across plasticity stages (“vertically”). In contrast, boundary height, jointly regulated by responsiveness and pliancy, showed phase-dependent, within-trial fluctuations and was modulated by plasticity in a more dynamic manner. The distinction between the two decision policy parameters has also been supported by previous behavioral modeling studies [49–51]. Across all these studies, the overall drift rate was found to increase steadily with learning, which was trial-independent [50] and was the most dramatic and consistent effect of training [49]. Conversely, the boundary height was a trial-dependent factor [50] and changed less systematically across training days and participants [51]. These studies, together with our analysis, highlight that drift rate primarily reflects experience-based action selection, whereas boundary height reflects strategic control for flexible adaptation across tasks and contexts.

Finally, the core conclusion of our work here is that dopamine-dependent corticostriatal plasticity exerts a strategic, time-resolved influence on decision formation by reshaping how evidence is accumulated across distinct phases of a single decision. Rather than simply changing overall choice biases or global decision thresholds, plasticity alters the relative engagement and timing of key CBGT populations, yielding a flexible and trajectory-dependent evidence accumulation process that can follow multiple paths before selection. To make these phase-dependent effects interpretable, we used the CLAW framework (cf. Figs 2 and 3A) as a descriptive lens to segment decision trajectories into biologically grounded regimes defined by activity patterns of the CBGT network. This view highlights that during a decision, learning-induced synaptic changes most strongly redirect network state, shifting the likelihood of distinct decision strategies and supporting within-decision adjustments rather than a single stereotyped accumulation profile (see also [32]).

Our CLAW analysis also connects CBGT circuit dynamics to recent low-dimensional descriptions of the decision-making process. Targeted dimensionality reduction (TDR) studies [52, 53] have shown that population activity can be organized around task-relevant axes in neural state space, with a dominant “choice axis” along which activity evolves during evidence integration. In this framework, position along the choice axis reflects progress towards making a selection, whereas momentary sensory or contextual inputs can transiently deflect trajectories into orthogonal dimensions, with collapse back toward the choice axis prior to decision time. Building on this perspective, our CLAW results provide a concrete instantiation of this conjectured framework within basal ganglia circuitry. In particular, the inner CLAW (deliberation) corresponds to the evidence integration dynamics, where trial trajectories unfold within a low-dimensional subspace before one action becomes stably dominant. The outer CLAW (commitment) corresponds to leaving this subspace, a divergence that locks into a specific decision. Importantly, our CLAW goes beyond the abstract axis-level description by identifying which CBGT populations actually drive these state transitions. In our model, learning-induced rise of choice-selective channel asymmetry, together with re-engagement of corticothalamic and direct pathways, provides a clear mechanism for driving trajectories into the outer branches, whereas indirect and pallidostriatal engagement supports maintenance of ongoing deliberation and regulates when a departure is permitted. In this way, corticostriatal plasticity can be understood as tuning both the evolution of trajectories during evidence integration (motion along a choice axis) and the conditions under which trajectories separate into choice-specific states (phase-dependent boundary control), yielding a mechanistic bridge between abstract manifold descriptions and biologically specified CBGT pathway interactions.

The phase-dependent control by plasticity leads directly to experimentally testable predictions about how learning could alter the detailed time course of basal ganglia dynamics during decisions. Building on our prior demonstration that CLAW analyses can extract meaningful decision dynamics from large-scale empirical recordings [32, 54], future work can assess whether trajectories at various stages of learning exhibit the same sequence and occupancy of states observed in our simulations (Fig 3A), and whether learning selectively amplifies or suppresses activity in the populations defining those states or zones. Similarly, even without direct knowledge of control ensembles (Figs 4, 5, and Supporting Information S3 Figure), recordings from key cell populations can be used to construct subnetwork proxies (e.g., Supporting Information S4 Figure) analogous to the control ensembles defined here. This process would enable the identification of key decision phases and quantification of phase-dependent learning effects. More broadly, future extensions can incorporate tonic dopamine contributions possibly related to urgency [55–57], changing reward contingencies [58–60], and richer choice settings with additional alternatives and more complex reward dynamics. Together, these analyses would determine how learning maximizes decision speed and accuracy across contexts while preserving the capacity for caution, deliberation, and mid-stream course corrections.

## Methods

### CBGT networks

The CBGT model implemented in this work is a biologically grounded spiking neural network consisting of ten distinct nuclei populations (Fig 1A): the cortical interneurons (CxI), the excitatory cortical neurons (Cx), the striatum, which includes the D1- and D2-expressing spiny projection neurons (dSPN and iSPN, respectively) and the fast-spiking interneurons (FSI), the globus pallidus external segment, which contains prototypical and arkypallidal neuron subtypes (GPeP and GPeA, respectively), the subthalamic nucleus (STN), the globus pallidus internal segment (GPi), and the thalamus (Th), derived from a sequence of previous studies [28, 29, 32, 36, 46]. In a simple two-choice task, we considered two separate populations for each neuronal type, corresponding to the left and right action channels, except for the CxI and FSI, which were shared across both channels. Decision trials were initiated by a constant stimulus input to the cortical neurons in both channels, driving an increase in stochastic cortical activity. This triggered the ramping activity across the striatal populations and impacted downstream basal ganglia and thalamic outputs. An action was declared to be selected when the instantaneous firing rate of the thalamic population in a specific channel reached 30 Hz before the other. Each neuron evolved according to a conductance-based integrate-and-fire-or-burst model. An example of firing rate activity across all populations, in the absence of synaptic plasticity, is shown in Fig 1B.

In our previous work [32], we used a genetic algorithm to generate 300 CBGT networks, each with a distinct configuration of synaptic connection weights (see also [31]). These networks were categorized into three equally sized groups—fast, intermediate, and slow—based on the mean decision time computed from trials sampled from each network simulation (in which no learning occurred). In the present study, we used the 100 slow networks, characterized by prolonged engagement of both deliberation and commitment, to investigate how learning impacted different phases within the decision process. Specifically, the corticostriatal projections to the dSPNs and iSPNs in the model were plastic. They were subject to a spike timing-dependent plasticity (STDP) rule, modulated by phasic dopamine signals elicited by decision outcomes [31, 59]. The critical learning signal emerged from a discrepancy between a received reward and the expected reward, known as the reward prediction error, which was encoded in the firing of dopaminergic neurons in the substantia nigra pars compacta (SNc). Here we assumed a deterministic reward contingency: trials selecting the left choice were consistently rewarded with 100% probability, whereas the right choice received no reward. Under these conditions, tuning on plasticity during simulations resulted in phasic dopamine release at the corticostriatal synapses and modified synaptic efficacy. This modulation influenced downstream activity in the basal ganglia and thalamic populations, thereby biasing subsequent decisions toward the rewarded direction. Complete details of the computational CBGT model, including implementation of the plasticity rule, can be found in [36].

### CLAW

In this work we replicated recent analysis of Circuit Logic Assessed via Walks (CLAW; see [32]) to compare the instantaneous neural dynamics underlying pre-learning and post-learning decisions, as depicted in Figs 2 and 3A. For each trial, the firing rate time series of each neural population was discretized into bins of Δ*t* = 10 ms and averaged within each bin. The average firing rate was then binarized based on whether it was above or below a pre-defined threshold, which was determined from the firing rate histogram aggregated across all simulated trials, as derived in [32]. We considered *N* = 10 key populations—dSPN, iSPN, GPi, GPeP, and Th for both left and right action channels—and we defined the set { *s*_*k*_ } in ℝ^*N*^ as the base-2 representations of all integers from 1 to 2^*N*^. Each *s*_*k*_ = [*σ*_*k*1_, *σ*_*k*2_, · · ·, *σ*_*kN*_], where *σ*_*kj*_ ∈ { 0, 1 } for all pairs (*k, j*), denotes the *k*-th unique state of the network. Full details of all states of the pre-learning scenario are presented in the Supporting Information S1 Table. In this way, we converted the CBGT firing rate time series to a sequence of states for every single trial. Using all of these state sequences, we computed the transition probabilities between every pair of possible states. Finally, we built a chain for the decision dynamics based on the state transitions, i.e., the CLAW, which extracted the most probable paths as decisions unfolded. Note that the state details listed in Figs 2B and 3B omit Th populations, since thalamic activity was shared across most states and thus contributed little to differentiating the displayed paths.

To reduce the impact of firing rate variability at individual states and to categorize key classes of decision trajectories, we further partitioned all CLAW states into a set of six unique zones (shaded blocks in Figs 2A and 3A), based on the transition probabilities and neural activity changes associated with transitions between states. States in each zone had common decision characteristics in terms of speed and choice. The partition also captured some occasional transitions that became apparent only when states were grouped into these broader zones. The whole CLAW analysis was conducted separately for pre-learning trials and for post-learning trials at each stage of learning. A comprehensive description of the approach, along with mechanistic insights into state and zone transitions, is provided in [32].

### Control ensembles

The DDM parameters and the CBGT activity for our 300 network configurations, before plasticity, were used to identify CBGT control ensembles through canonical correlation analysis (CCA), as was done in prior work [29, 31, 32]. The outcomes of trials from our simulated CBGT networks were mapped to the static DDM by fitting their decision times and choices, using the Hierarchical Sequential Sampling Modeling (`HSSM`) toolbox implemented in Python, and we obtained the corresponding configuration of static DDM parameters (boundary height *a*, drift rate *v*, onset time *tr*, and starting bias *z*) for each network. CCA then found pairs of linear combinations, i.e., canonical variates, from the space of CBGT activity elements (including channel-summed activity and between-channel differences, 18 elements in total) and the space of DDM parameters, that were maximally correlated. From these, we selected three key components that captured the dominant shared variance between the two datasets, as illustrated in Fig 4. The corresponding canonical loadings, which quantify the contribution of each CBGT element and DDM parameter to their respective canonical variates, are listed in the Supporting Information S2 Table.

Using control ensembles, we converted the time-binned series of CBGT firing rates to a discretized time series of individual control ensemble engagement. Following the method described in [32] (see also [31]), for each trial we computed Δ*F*_*k*_ ∈ ℝ^18^, representing the difference in average firing rates of CBGT elements between time bin *k* and cue onset. This difference was then projected onto the three control ensemble components, via 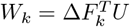 where the columns of matrix *U* contain the canonical loadings related to CBGT elements. As a result, each vector *W*_*k*_ comprises three elements, representing changes in the corresponding control ensemble engagement as the trial evolved to the *k*-th bin. Note that Δ*F*_0_ was always set to zero as a baseline (at time 0 on the left end of the x-axis), representing the initial state of firing rates before any changes in the decision process occurred. Finally, we collected the discretized time series of each element of *W*_*k*_ across trials to examine the overall evolution of control ensemble engagement, as shown in Figs 5 and S3 Figure. In the last step, to link these dynamics to DDM parameters, we further projected *W*_*k*_ to the DDM space via *P*_*k*_ = *W*_*k*_*V* ^*T*^ where the loadings related to DDM parameters comprise the columns of *V*. This allowed us to estimate the average change in drift rate and boundary height as decisions unfolded, as displayed in Fig 6. A justification of the validity of using CCA to infer dynamic, within-trial changes in decision policies can be found in [32].

### Learning effect

We employed a mixed-effects regression model to examine whether learning significantly affected the temporal evolution of control ensemble engagement (Figs 5 and S3 Figure) and DDM parameters (Fig 6). The model was implemented using the `MixedLM` function from the `statsmodels` library in Python. In each decision class comprising two types of trajectories, we treated learning (quantified by the number of training trials) and trajectory type (categorical) as fixed effects, while network identity was modeled as a random effect to account for variability across network configurations. Both main effects and interaction terms among time, learning, and trajectory type were included in the model. Given the possibility of nonlinearity in the time series, particularly during the commitment phase, we tested both linear and quadratic models (including time and time^2^ terms) and selected the one with the lowest Bayesian information criterion (BIC) score. Since the analysis was conducted independently for each of the four decision classes, we corrected for multiple comparisons using a Bonferroni adjustment, setting the significance threshold at *α* = 0.05*/*4 = 0.0125.

Our primary focus was on interaction terms involving time and learning, time and trajectory type, and the three-way interaction among these variables (including the quadratic counterparts if the quadratic model was optimal). These terms allowed us to assess whether learning and trajectory type significantly modulated the overall temporal profiles of control ensemble engagement and DDM parameters. In cases where significant effects were observed over the full trajectory, we performed a follow-up pointwise analysis at individual time points to determine when during the decision process these effects emerged. A standard significance threshold (*α* = 0.05) was used for this step. Model complexity at each time point was adapted based on the presence of significant main effects and/or interactions observed in the preceding full-trajectory analysis. The horizontal bars above each panel in Figs 5, 6 and S3 Figure indicate the time intervals during which learning exerted a significant effect for the corresponding trajectories.

### Dynamic DDM fitting

We developed a dynamic DDM fitting approach to capture how the decision policy parameters, particularly drift rate *v* and boundary height *a*, evolved as during the evidence accumulation process. This approach was built upon the static DDMs but integrated time-varying information derived from CBGT network dynamics. First, for each trajectory type at each learning stage, we fit the decision times and choices of the associated trials with the static DDM, implemented via the `HSSM` toolbox. This fit yielded a single set of DDM parameters, denoted as (*v*_static_, *a*_static_, *tr*_static_, *z*_static_), based on the assumption of a time-homogeneous evidence accumulation process over the entire decision period. Next, we calculated the change in the CBGT activity elements (described in Control ensembles) as trial trajectories proceeded through differet CLAW zones. For instance, for *p* trials that followed the path I→II→III, we computed Δ*F*_12_, Δ*F*_23_ ∈ ℝ^*p×*18^, representing the differences from zones I→II and II→III, respectively, and also recorded the average time spent in each zone, denoted as *t*_1_, *t*_2_, and *t*_3_. These activity differences were projected onto the DDM parameter space using the canonical loadings from the previously identified control ensembles. As a result, we obtained the percentage changes in drift rate and boundary height driven by each zone transition. For example, suppose that the projection indicated an average decrease of *x*% in *a* from zone I to II, and a further decrease of *y*% from zone II to III. By using these relative changes, we then estimated actual boundary height in each zone. The estimates were constrained such that the time-weighted average of the dynamic boundary heights matched the static DDM fit:

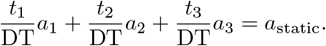

Substituting the relationships

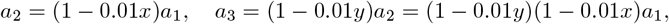

we solved for *a*_1_ and obtained the corresponding values of *a*_2_ and *a*_3_. The same calculation was applied to derive dynamic values for the drift rate *v*. In this way, we were able to find the dynamic (*v*_*i*_, *a*_*i*_) values as decision trajectories flowed through different CLAW zones.

To validate our approach, we simulated a piecewise DDM in which the accumulated evidence variable *θ* evolved according to the estimated dynamic parameters. Specifically,

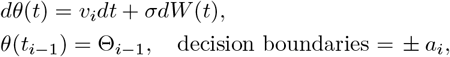

where Θ_*i*−1_ is the ending position of the previous phase,

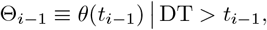

i.e., the value at the phase transition conditioned on there being no decision (boundary-crossing) before time *t*_*i*−1_. Because onset time *tr* and starting bias *z* defined the initial conditions, we used their static values, i.e.,

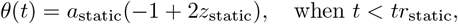

and simulations began with (*v*_1_, *a*_1_). A key challenge was to determine the appropriate time at which the model should transition to (*v*_2_, *a*_2_) and then to (*v*_3_, *a*_3_), since the simulated DDM did not have specific zones or phases. To address this, we sampled transition times stochastically from a categorical distribution over zone indices, weighted by the relative time spent in each zone as reported from the CBGT trajectories. If transition to zone II occurred before *θ* reached the boundaries *±a*_1_, the simulation continued with parameters (*v*_2_, *a*_2_); otherwise, the trial was discarded. The same logic was applied for transitions to zone III. Note that in cases where *θ* remained below *a*_2_ but exceeded *a*_3_ at the transition point, the immediate boundary collapse to *a*_3_ was interpreted as a decision, and the trial was included. Across all simulations, the diffusion noise level *σ* was fixed at 1, consistent with the implementation via `HSSM`.

This procedure ensured that only those paths consistent with the timing and parameter changes inferred from CBGT data were retained. Examples of such paths are illustrated in Fig 9A, contrasted with those simulated by the static version in Fig 9B. The resulting decision time distributions from both dynamic and static simulations, along with the simulated distribution derived from the CBGT data, are also shown in the corresponding bottom panels. The dynamic model achieved a significantly lower BIC compared to the static one and produced mean decision times more closely aligned with the observed CBGT data. This improved performance implied that by allowing drift rate and boundary height to change across decision phases, our dynamic fitting approach provided a more realistic representation of the decision-making process, effectively reflecting the evolving dynamics of the underlying CBGT circuit. Following this approach for each decision type at each plasticity stage, we summarized in Fig 8 the corresponding decision policy trajectory in the (|*v*|, *a*)-space.

**Fig 9.**
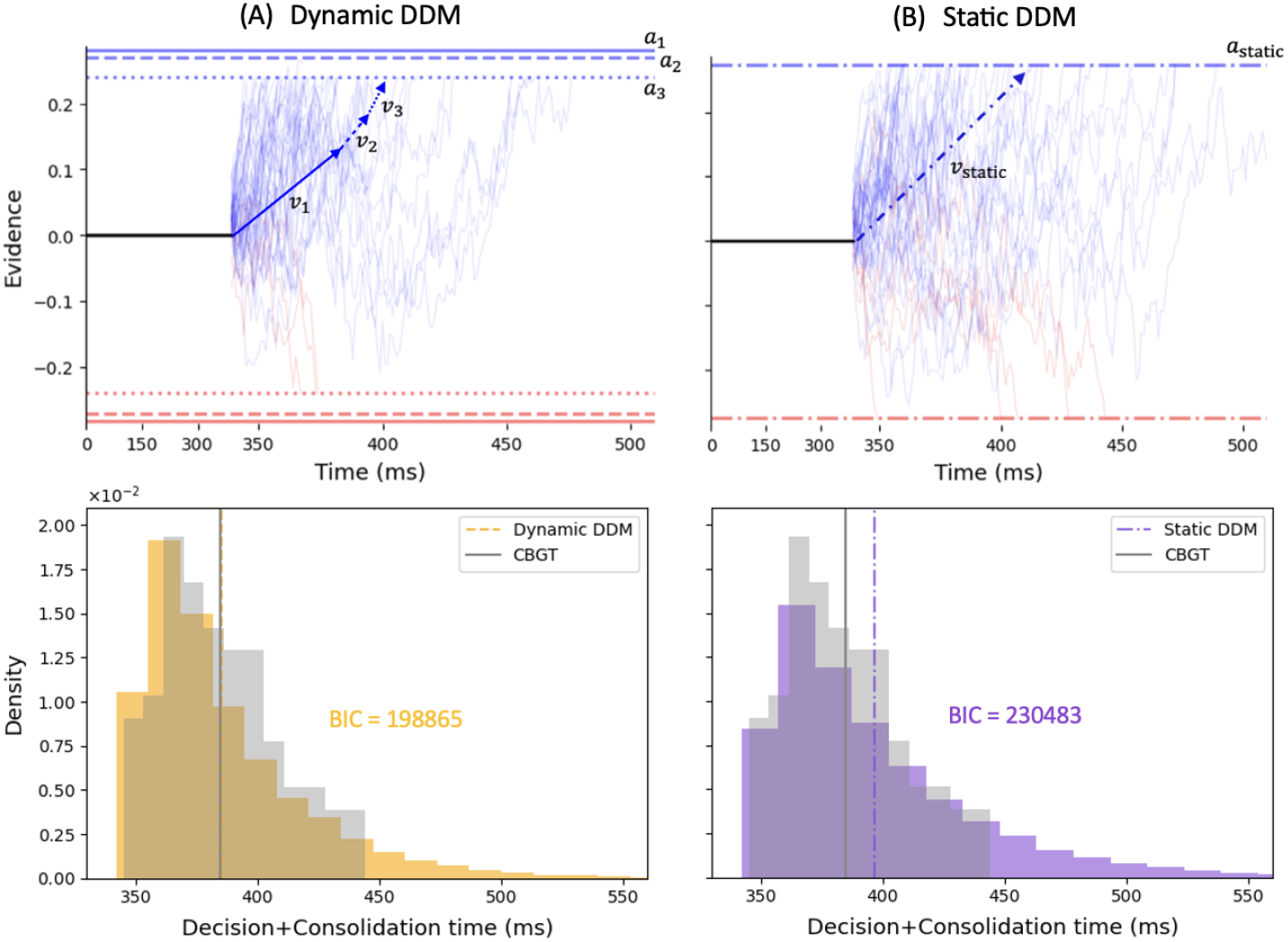
Comparison between dynamic and static DDM simulations. The DDM parameters were estimated from fitting the chocies and decision+consolidation times (i.e., pink+grey regions in Fig 1B, where consolidation times were sampled from *N* (250, 1.5)) of pre-learning trials within trajectory type I→II→III. The dynamic model (A) allowed drift rate *v* and boundary height *a* to vary during evidence accumulation, with *a* decreasing sequentially from *a*_1_ (solid) to *a*_2_ (dashed) to *a*_3_ (dotted), and *v* increasing from *v*_1_ to *v*_2_ to *v*_3_. The static model (B) assumed fixed parameters, *a*_static_ and *v*_static_ (dash-dotted), throughout evidence accumulation. Bottom panels show the corresponding decision+consolidation time distribution (yellow for dynamic and purple for static), compared with the simulated CBGT data (grey). The mean of each distribution is marked by a vertical line of the corresponding color. The dynamic DDM model yielded a significantly lower BIC and a mean decision+consolidation time that more closely matched the CBGT data.

## Supporting information

**S1 Table. CLAW state details**. The table in Fig 2B includes the details of the left-related and neutral states in the upper half of the CLAW, while this table provides the information for all CLAW states, with each pair of left- and right-related states showing symmetry up to the swap of certain L and R channel binary values.

**S2 Table. Control ensemble loadings**. Figure 4 illustrates the relation between the activity of each CBGT population (either summed across channels or as between-channel differences) and the DDM parameters, obtained in the first three components of the CCA. This table provides the corresponding loadings for each element of CBGT activity and DDM.

**S3 Figure. Control ensemble engagement for each decision type**. As an expanded companion to Fig 5, this figure shows control ensemble time courses for each decision type across learning stages. Each point along the trace represents the percent change in the control ensemble engagement relative to cue onset. Each decision class included two types of trajectories, shown in blue and red. Trials of each type were aligned to cue onset (time 0 on the left end of the x-axis) and decision time (time 0 on the right). Trace hues indicate learning progression, from pre-learning (light) to early learning (medium) to late learning (dark). Shaded horizontal bars above each panel indicate time bins in which the learning effect on the corresponding control ensemble was statistically significant, with bar colors matching the associated type.

**S4 Figure. Key nuclei firing rates pre- and post-learning**. This figure compares choice-selective (differential) and global (overall) firing activity in core cell populations across three CBGT pathways before and after learning, providing testable predictions for future experiments. Firing rates were averaged over trials sampled from four left-related trajectories: I → II, I → II → V, I → II → III, and I → III. Trials at each learning stage were aligned to cue onset (time 0) but not to decision time to highlight the difference in actual decision times between pre-learning (light blue) and post-learning (dark blue). Time series from each trial were normalized to the mean decision time within each condition—130 ms for pre-learning and 100 ms for post-learning. Dotted lines in grey regions correspond to the launching phase we defined (first 50 ms after cue onset); dashed lines correspond to the intermediate period; solid lines correspond to the committed phase (final 30 ms prior to decision). Shadow areas around the mean firing rate traces represent the 95% confidence interval across trials.

## Acknowledgments

We thank Jyotika Bahuguna, Cati Vich Llompart, and Eric Yttri for helpful comments on this work. This work was supported by NIH award R01DA059993 as part of the CRCNS program. JER is also partly supported by NIH award R01NS125814, also part of the CRCNS program.

## References

1. Humphries MD, Stewart RD, Gurney KN. A physiologically plausible model of action selection and oscillatory activity in the basal ganglia. Journal of Neuroscience. 2006;26(50):12921–12942.

2. Bogacz R, Gurney K. The basal ganglia and cortex implement optimal decision making between alternative actions. Neural computation. 2007;19(2):442–477.

3. Doya K. Modulators of decision making. Nature neuroscience. 2008;11(4):410–416.

4. Chakravarthy VS, Joseph D, Bapi RS. What do the basal ganglia do? A modeling perspective. Biological cybernetics. 2010;103(3):237–253.

5. Dunovan K, Verstynen T. Believer-skeptic meets actor-critic: rethinking the role of basal ganglia pathways during decision-making and reinforcement learning. Frontiers in neuroscience. 2016;10:106.

6. Schultz W, Apicella P, Scarnati E, Ljungberg T. Neuronal activity in monkey ventral striatum related to the expectation of reward. Journal of neuroscience. 1992;12(12):4595–4610.

7. Mirenowicz J, Schultz W. Importance of unpredictability for reward responses in primate dopamine neurons. Journal of neurophysiology. 1994;72(2):1024–1027.

8. Hollerman JR, Schultz W. Dopamine neurons report an error in the temporal prediction of reward during learning. Nature neuroscience. 1998;1(4):304–309.

9. Reynolds JN, Hyland BI, Wickens JR. A cellular mechanism of reward-related learning. Nature. 2001;413(6851):67–70.

10. Roesch MR, Calu DJ, Schoenbaum G. Dopamine neurons encode the better option in rats deciding between differently delayed or sized rewards. Nature neuroscience. 2007;10(12):1615–1624.

11. Albin RL, Young AB, Penney JB. The functional anatomy of basal ganglia disorders. Trends in neurosciences. 1989;12(10):366–375.

12. DeLong MR. Primate models of movement disorders of basal ganglia origin. Trends in neurosciences. 1990;13(7):281–285.

13. Bariselli S, Fobbs W, Creed M, Kravitz A. A competitive model for striatal action selection. Brain research. 2019;1713:70–79.

14. Alexander GE, DeLong MR, Strick PL. Parallel organization of functionally segregated circuits linking basal ganglia and cortex. Annual review of neuroscience. 1986;9(1):357–381.

15. Alexander GE, Crutcher MD. Functional architecture of basal ganglia circuits: neural substrates of parallel processing. Trends in neurosciences. 1990;13(7):266–271.

16. Middleton FA, Strick PL. Basal ganglia and cerebellar loops: motor and cognitive circuits. Brain research reviews. 2000;31(2-3):236–250.

17. Mink JW. The basal ganglia: focused selection and inhibition of competing motor programs. Progress in neurobiology. 1996;50(4):381–425.

18. Mallet N, Micklem BR, Henny P, Brown MT, Williams C, Bolam JP, et al. Dichotomous organization of the external globus pallidus. Neuron. 2012;74(6):1075–1086.

19. Giossi C, Rubin JE, Gittis A, Verstynen T, Vich C. Rethinking the external globus pallidus and information flow in cortico-basal ganglia-thalamic circuits. European Journal of Neuroscience. 2024;60(9):6129–6144.

20. Mallet N, Pogosyan A, Márton LF, Bolam JP, Brown P, Magill PJ. Parkinsonian beta oscillations in the external globus pallidus and their relationship with subthalamic nucleus activity. Journal of neuroscience. 2008;28(52):14245–14258.

21. Nevado-Holgado AJ, Mallet N, Magill PJ, Bogacz R. Effective connectivity of the subthalamic nucleus–globus pallidus network during Parkinsonian oscillations. The Journal of physiology. 2014;592(7):1429–1455.

22. Ketzef M, Silberberg G. Differential synaptic input to external globus pallidus neuronal subpopulations in vivo. Neuron. 2021;109(3):516–529.

23. Mallet N, Schmidt R, Leventhal D, Chen F, Amer N, Boraud T, et al. Arkypallidal cells send a stop signal to striatum. Neuron. 2016;89(2):308–316.

24. Aristieta A, Barresi M, Lindi SA, Barriere G, Courtand G, de La Crompe B, et al. A disynaptic circuit in the globus pallidus controls locomotion inhibition. Current Biology. 2021;31(4):707–721.

25. Giossi C, Bahuguna J, Rubin JE, Verstynen T, Vich C. Arkypallidal neurons in the external globus pallidus can mediate inhibitory control by altering competition in the striatum. Proceedings of the National Academy of Sciences. 2024;121(47):e2408505121.

26. Simen P, Cohen JD, Holmes P. Rapid decision threshold modulation by reward rate in a neural network. Neural networks. 2006;19(8):1013–1026.

27. Ratcliff R, Frank MJ. Reinforcement-based decision making in corticostriatal circuits: mutual constraints by neurocomputational and diffusion models. Neural computation. 2012;24(5):1186–1229.

28. Dunovan K, Vich C, Clapp M, Verstynen T, Rubin J. Reward-driven changes in striatal pathway competition shape evidence evaluation in decision-making. PLoS computational biology. 2019;15(5):e1006998.

29. Vich C, Clapp M, Rubin JE, Verstynen T. Identifying control ensembles for information processing within the cortico-basal ganglia-thalamic circuit. PLOS Computational Biology. 2022;18(6):e1010255.

30. Ratcliff R. A theory of memory retrieval. Psychological review. 1978;85(2):59.

31. Bahuguna J, Verstynen T, Rubin JE. How cortico-basal ganglia-thalamic subnetworks can shift decision policies to maximize reward rate. bioRxiv. 2025; p. 2024–05.

32. Yu Z, Verstynen T, Rubin JE. How the dynamic interplay of cortico-basal ganglia-thalamic pathways shapes the time course of deliberation and commitment. bioRxiv. 2025; p. 2025–03.

33. Srivastava V, Feng SF, Cohen JD, Leonard NE, Shenhav A. A martingale analysis of first passage times of time-dependent Wiener diffusion models. Journal of mathematical psychology. 2017;77:94–110.

34. Holmes WR, Trueblood JS. Bayesian analysis of the piecewise diffusion decision model. Behavior research methods. 2018;50(2):730–743.

35. Palestro JJ, Weichart E, Sederberg PB, Turner BM. Some task demands induce collapsing bounds: Evidence from a behavioral analysis. Psychonomic bulletin & review. 2018;25(4):1225–1248.

36. Clapp M, Bahuguna J, Giossi C, Rubin JE, Verstynen T, Vich C. CBGTPy: An extensible cortico-basal ganglia-thalamic framework for modeling biological decision making. PloS one. 2025;20(1):e0310367.

37. Young PA, Waller O, Ball K, Williams CC, Nashmi R. Phasic stimulation of dopaminergic neurons of the lateral substantia nigra increases open field exploratory behaviour and reduces habituation over time. Neuroscience. 2024;551:276–289.

38. Chakroun K, Mathar D, Wiehler A, Ganzer F, Peters J. Dopaminergic modulation of the exploration/exploitation trade-off in human decision-making. Elife. 2020;9:e51260.

39. Bakhurin KI, Li X, Friedman AD, Lusk NA, Watson GD, Kim N, et al. Opponent regulation of action performance and timing by striatonigral and striatopallidal pathways. Elife. 2020;9:e54831.

40. Yartsev MM, Hanks TD, Yoon AM, Brody CD. Causal contribution and dynamical encoding in the striatum during evidence accumulation. Elife. 2018;7:e34929.

41. Dhawale AK, Smith MA, Ölveczky BP. The role of variability in motor learning. Annual review of neuroscience. 2017;40(1):479–498.

42. Dhawale AK, Miyamoto YR, Smith MA, Ölveczky BP. Adaptive regulation of motor variability. Current Biology. 2019;29(21):3551–3562.

43. Wu HG, Miyamoto YR, Castro LNG, Ölveczky BP, Smith MA. Temporal structure of motor variability is dynamically regulated and predicts motor learning ability. Nature neuroscience. 2014;17(2):312–321.

44. Cui G, Jun SB, Jin X, Pham MD, Vogel SS, Lovinger DM, et al. Concurrent activation of striatal direct and indirect pathways during action initiation. Nature. 2013;494(7436):238–242.

45. Kobayashi K, Fukabori R, Nishizawa K. Neural circuit mechanism for learning dependent on dopamine transmission: roles of striatal direct and indirect pathways in sensory discrimination. Advances in Pharmacology. 2013;68:143–153.

46. Wei W, Rubin JE, Wang XJ. Role of the indirect pathway of the basal ganglia in perceptual decision making. Journal of Neuroscience. 2015;35(9):4052–4064.

47. Li H, Jin X. Multiple dynamic interactions from basal ganglia direct and indirect pathways mediate action selection. Elife. 2023;12:RP87644.

48. Kim SY, Lim W. Quantifying harmony between direct and indirect pathways in the basal ganglia: healthy and Parkinsonian states. Cognitive Neurodynamics. 2024;18(5):2809–2829.

49. Eckhoff P, Holmes P, Law C, Connolly P, Gold J. On diffusion processes with variable drift rates as models for decision making during learning. New Journal of Physics. 2008;10(1):015006.

50. Pedersen ML, Frank MJ, Biele G. The drift diffusion model as the choice rule in reinforcement learning. Psychonomic bulletin & review. 2017;24(4):1234–1251.

51. Cochrane A, Sims CR, Bejjanki VR, Green CS, Bavelier D. Multiple timescales of learning indicated by changes in evidence-accumulation processes during perceptual decision-making. npj Science of Learning. 2023;8(1):19.

52. Mante V, Sussillo D, Shenoy KV, Newsome WT. Context-dependent computation by recurrent dynamics in prefrontal cortex. nature. 2013;503(7474):78–84.

53. Pagan M, Tang VD, Aoi MC, Pillow JW, Mante V, Sussillo D, et al. Individual variability of neural computations underlying flexible decisions. Nature. 2025;639(8054):421–429.

54. Steinmetz NA, Zatka-Haas P, Carandini M, Harris KD. Distributed coding of choice, action and engagement across the mouse brain. Nature. 2019;576(7786):266–273.

55. Beeler JA, Daw N, Frazier CR, Zhuang X. Tonic dopamine modulates exploitation of reward learning. Frontiers in behavioral neuroscience. 2010;4:170.

56. Humphries MD, Khamassi M, Gurney K. Dopaminergic control of the exploration-exploitation trade-off via the basal ganglia. Frontiers in neuroscience. 2012;6:16922.

57. Gilbertson T, Steele D. Tonic dopamine, uncertainty and basal ganglia action selection. Neuroscience. 2021;466:109–124.

58. Frank MJ, Gagne C, Nyhus E, Masters S, Wiecki TV, Cavanagh JF, et al. fMRI and EEG predictors of dynamic decision parameters during human reinforcement learning. Journal of Neuroscience. 2015;35(2):485–494.

59. Vich C, Dunovan K, Verstynen T, Rubin J. Corticostriatal synaptic weight evolution in a two-alternative forced choice task: a computational study. Communications in Nonlinear Science and Numerical Simulation. 2020;82:105048.

60. Beron CC, Neufeld SQ, Linderman SW, Sabatini BL. Mice exhibit stochastic and efficient action switching during probabilistic decision making. Proceedings of the National Academy of Sciences. 2022;119(15):e2113961119.

